# Eco-evolutionary dynamics of planktonic calcifying communities under ocean acidification

**DOI:** 10.64898/2026.03.01.708833

**Authors:** Théo Villain, Nicolas Loeuille

## Abstract

Increasing emissions of CO_2_ into the atmosphere are causing ocean acidification, threatening calcifying organisms. In this study, we model the physiological responses of coccolithophorids to acidification to understand the ecological and evolutionary outcomes of a system in interaction with zooplankton. Assuming a trade-off between growth and protection against grazing, we show that calcification has bivalent effects on transfers between two trophic levels and that acidity can strongly alter energy transfers. Taking into account the evolution of calcifying phenotypes in response to acidification, we show that the system outcome contrasts with previous results. While the effect of evolution depends on how calcification affects grazing, it nevertheless follows that acidification leads to a decrease in calcifying capacity. This evolutionary decrease may be progressive, but can also lead to tipping points where abrupt shifts may occur. Such a counter-selection of calcification in turn affects ecosystem functioning, enhancing energy transfers within the system and modifying carbon fluxes. We discuss how such eco-evolutionary changes may impact food webs integrity, carbon sequestration into the deep ocean and therefore endanger the carbon pump stability.

## Introduction

Ocean acidification (OA) is a major threat for the functioning of marine ecosystems (Doney et al., 2009; Fabry et al., 2008; Kleypas and Yates, 2009). The ocean is a major carbon reservoir on Earth and plays an essential role in regulating the global carbon cycle, absorbing an estimated 22 million tons of carbon dioxide (CO_2_) per day (Feely et al., 2008). OA is therefore caused by the diffusion into marine waters of a quarter of the atmospheric CO_2_ (Sabine et al., 2004) currently enhanced by ever-increasing fossil fuel combustion (Houghton et al., 2001). Dissolved CO_2_ changes seawater pH and the reaction equilibrium of ocean carbonates according to the system CO_2_ + H_2_O = H^+^ + HCO_3_ ^-^= 2H^+^ + CO_3_ ^2-^ (Jansen et al., 2002). Buffered by a preferential association of protons with CO_3_ ^2-^, OA leads to a decrease in its concentration while it is a major substrate for calcite or aragonite precipitation (CaCO_3_). Most pessimistic IPCC scenarios RCP8.5 project a rise in atmospheric carbon dioxide (440 to 1000 ppm) in 2100 causing marine pH to decrease from 8.15 to 7.79 on average, associated with rising seawater CO_2_ concentrations (15 µmol.L^−1^ to 35 µmol.L^−1^).

These projections suggest the decline of carbonate CO_3_ ^2-^ concentrations from 180 µmol.L^−1^ to 90 µmol.L^−1^ in the same period (Riahi et al., 2011). Under this scenario, CO_2_ concentrations may reach [CO_2_] = 70 µmol.L^−1^ by 2300 (Caldeira and Wickett, 2003). Because Ω (the saturation state of seawater in calcite CaCO_3_) thermodynamically controls calcite precipitation or dissolution, supersaturated waters (Ω > 1) favors precipitation whereas undersaturation (Ω < 1) leads to calcite dissolution. Between 200 and 3500 m depth, the CaCO_3_ saturation horizon (Ω = 1) is currently rising towards the surface which constrains areas where calcification could thermodynamically occur. Combined with direct effects on pH, decreasing CO_3_ ^2-^ concentrations have caused the horizon to rise from 80 to 200 m since the beginning of the industrial area (Feely et al., 2004). With a rate of 4 m/year (Bijma et al., 2013), it could reach the surface of the Southern Ocean by 2100 (Intergovernmental panel on climate change, 2007). Next to these global trends, local variations may largely vary in amplitude and in time. For instance, while subtropical surface waters are generally CO_2_-rich in summer, Arctic Ocean contains lowest concentration during July-August (Orr et al., 2022) and combined with seasonal variation of temperature and winds it is certainly responsible for carbonate ion (CO_3_ ^2-^) strong depletion in winter (McNeil and Matear, 2008).

These changes in ocean chemistry directly affect the physiology of marine organisms, particularly those precipitating carbonates to produce thallus, skeletons, shells, coralline structures or protections. The current trend towards carbonate ion undersaturation due to OA strongly affects CaCO_3_ solubility and thus calcifying capacities of organisms (Pandolfi et al., 2011). Because many of them are pelagic (such as coccolithophorids) or benthic (coralline algae, corals) primary producers and since calcification provides protection against predators (Monteiro et al., 2016), OA effects on their physiology can cascade through the entire trophic chain, altering bottom-up regulations within the system and drastically changing community structures (Dutkiewicz et al., 2015; Meakin and Wyman, 2011; Taucher et al., 2017) differently according to latitudes (Hoppe et al., 2018). Some studies reveal that variations in aragonite saturation states induce changes from coralline algae-dominated systems to organic algae-dominated in mesocosm manipulations and might strongly alter benthic community assemblages (Kuffner et al., 2008). Ecosystem engineering by calcifying organisms such as corals (Erwin, 2008) is seriously threatened by OA among other climatic changes (Anthony et al., 2008; Hoegh-Guldberg et al., 2017; Pandolfi et al., 2011) which could have strong indirect effects on communities by alteration of the ecological functions corals provide (niche construction, food supply) (Hoegh-Guldberg et al., 2017; Kleypas and Yates, 2009).

Coccolithophorids (calcifying phytoplankton, e.g. Brownlee and Taylor, 2004) alone are responsible for 1-10% of net primary planetary productivity (Poulton et al., 2007), which represents about 21 Pg of carbon fixed per year (Field et al., 1998). These haptophytes account for more than 80% of the calcification produced on an oceanic scale (Milliman, 1993; Westbroek et al., 1989). These photosynthetic species are distributed in the euphotic zone and by converting inorganic carbon into organic matter, they allow the transfer of carbon to the bottom of the ocean (organic carbon pump). This is of primary importance in setting up the vertical CO_2_ gradient and more generally in regulating the marine carbon cycle and its exchange with the atmosphere (Holligan and Robertson, 1996). Calcification, on the other hand, causes a release of carbon on the surface (H_2_O + CO_2_ + CaCO_3_ = 2HCO_3_ ^-^+ Ca^2+^) but the rain of carbonate matter from these organisms towards ocean floor allows carbon storage in the sediments and its dissolution is also a carbon sink (carbonate counter pump phenomenon, Denman et al., 2007). By reducing the transfer up of carbon in food webs and increasing its sinking rates into the sea floor, calcification is strongly associated with the marine carbon cycle. Disrupting the physiology of these organisms by OA can have global consequences on oceanic trophic chains, marine chemistry and the global carbon pump.

The effects of OA on mass bleaching of many coral reefs have already been documented in numerous papers (e.g. Gattuso, 1998; Pandolfi et al., 2011) and the number of experiments examining these effects in different species of calcifying phytoplankton has been increasing exponentially over the last two decades (e.g. Kroeker et al., 2010; Meyer and Riebesell, 2015; Riebesell et al., 2000; Schlüter et al., 2016, 2014). Short-term physiological studies suggest that OA tends to decrease the ability to calcify and to produce energy through photosynthesis. Nonetheless most studies focus on one or a few species. Therefore, they cannot account for indirect ecosystemic effects (e.g. variations in grazing pressures) nor can they assess the implications of OA for the functioning of marine food webs. Given the importance of calcification in herbivory resistance (e.g. Monteiro et al., 2016), this scale should be considered. Indeed, coccoliths (calcareous structures secreted by coccolithophorids), among other functions, defend the cell against herbivores and limit the top-down control that phytoplankton undergo. The counter selection of calcification by acidification could then modify the vertical energy flow within the systems.

OA not only affects calcifying capacities but also constrains their evolutionary dynamics due to direct (changes in pH) and indirect effects (changes in the ecological context, eg, decrease in herbivory pressure) (Meyer and Riebesell, 2015). Experimental evidence suggests that such an evolution can be rapid and lead to large changes in resilience and productivity (Schlüter et al., 2014). Such evolutionary experiments however focus on the evolving species, and do not include variations of the surrounding communities, including grazers. Using an eco-evolutionary model that links physiological aspects of calcification and protection against grazing, we investigate whether i) coccolithophorids calcifying abilities change under OA, ii) under what conditions eco-evolutionary dynamics enable community maintenance and iii) how rising atmospheric CO_2_ could affect community structure, diversity and potential carbon fluxes. We predict that acidification will ultimately counterselects calcification ability, thereby increasing available energy for higher trophic levels and decreasing sinking carbon fluxes.

## 1. Methods

### 1.1 Definition of x, the energy investment in calcification

CO_2_ diffuses into the cell in dissolved form or combined with water. The coccolithophorid then distributes the captured inorganic carbon between photosynthesis (the energy source of all the other functions) and calcification. The resulting chemical energy (carbohydrate) is distributed between calcification and other functions within cells. We take it into account substrate partitioning (CO_2_ for photosynthesis and HCO3-for calcification (Gafar et al., 2018)) but do not model the resulting cytosolic competition. The energy from photosynthesis is partitioned into a fraction *x* allocated to calcification, *(1-x)* being allocated to growth. This hypothesis defines a trade-off according to *x* between growth and calcification. As protection against enemies is one of the putative selective advantages of calcification (Monteiro et al., 2016), we assume that high calcifiers (ie, high *x*) leads to lower herbivory pressure. On the other hand, low calcifiers have more energy to perform survival, growth and reproduction. This type of defense-growth trade-off is not restricted to our system, but is often observed in ecology, for instance in terrestrial plants (Herms and Mattson, 1992). This trait *x* is therefore an energetic compromise on a physiological scale, but it also has ecological consequences for the dynamics of phytoplankton and zooplankton populations. Our aim is here to uncover how this trait evolves in response to OA. Numerous experimental studies have tackled this by calculating a PIC/POC ratio (see the meta-analyses by Kroeker et al., 2010; Meyer and Riebesell, 2015) and show that investment in calcification clearly tends to decrease at lower pH. These monospecific experiments however focus on direct population responses and do not take into account changes in the ecological context.

### 1.2 The ecological model

Because we intend to account for both physiological constraints and for evolutionary dynamics (see below), we here keep the ecological dynamics simple. Trophic interactions between phytoplankton investing *x* in calcification and zooplankton at a given CO_2_ concentration are therefore modeled using a simple Lotka-Volterra model:

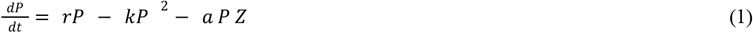

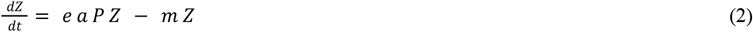

*P* and *Z* are the respective densities (per mL) of calcifying phytoplankton and zooplankton, *r* the intrinsic growth rate of phytoplankton, *k* the competition coefficient of phytoplankton, *a* the attack rate of zooplankton on phytoplankton, *e* the conversion factor of zooplankton and *m* the mortality of zooplankton.

#### 1.2.1 Intrinsic growth rate

Energy from photosynthesis is modeled by a Monod function **π**_p_ (eq. 4) (e.g. Paul and Bach, 2020; Riebesell et al., 1993) based on data from Sett et al., 2014. This experimental study measures the growth and calcification rates of *Emiliania huxleyi* exposed to different CO_2_ concentrations and serves as the basis for our modeling. The intrinsic mortality of phytoplankton depends on *x*. The higher *x*, the lower the relative amount of energy available to maintain cellular homeostasis. The intrinsic growth rate of phytoplankton is therefore based on the proportion of energy left for the growth and survival of the organism minus the intrinsic mortality of the cell (see Table 2 for a description of the parameters used):

**Table 1.**
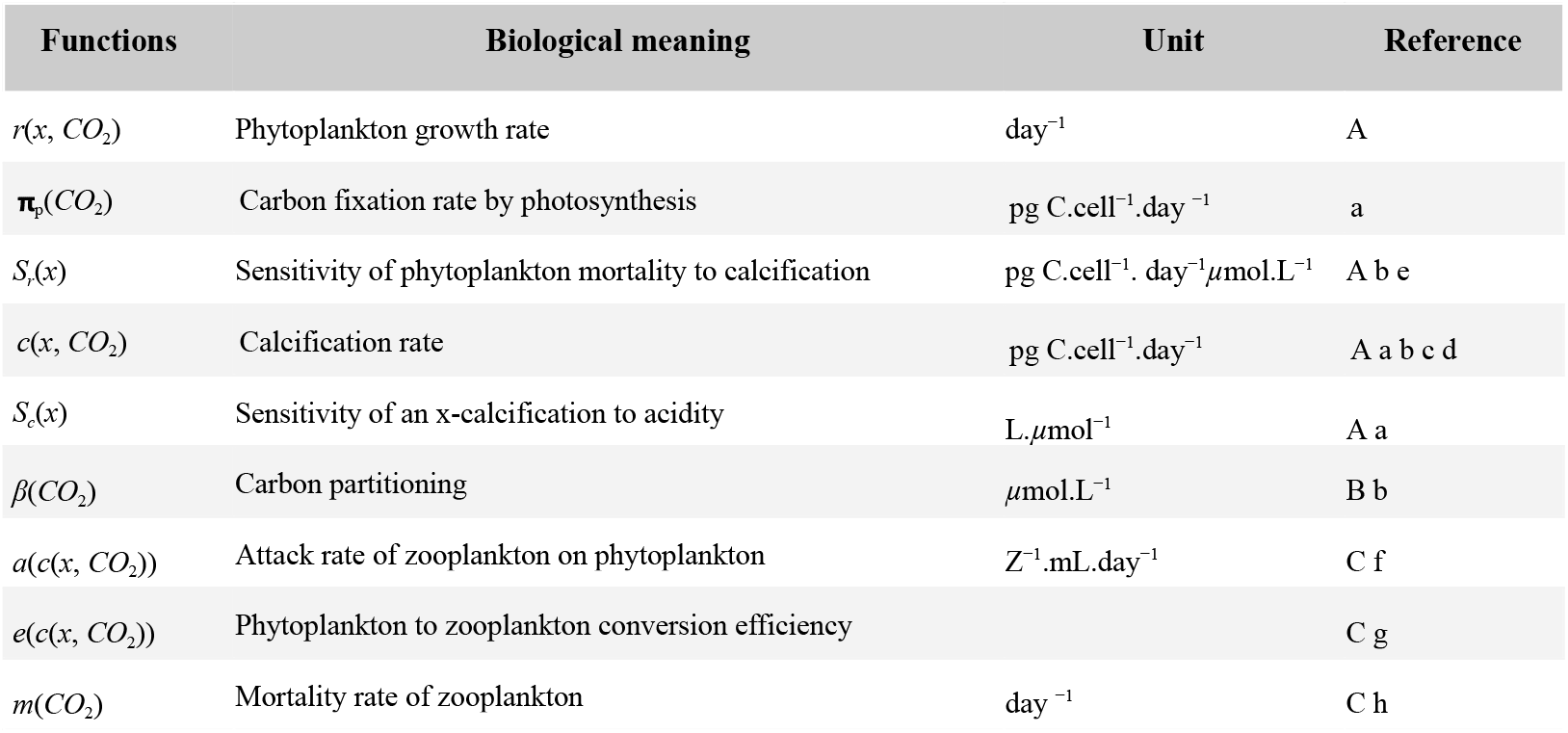
Model Functions **a** : Sett et al., 2014, **b** : Bach et al., 2015, **c** :Bach et al., 2012, **d** : Bach et al., 2011, **e** : Quéré et al., 2005, **f** :Harvey et al., 2015, **g** : Haunost et al., 2021, **h** : Cripps et al., 2014; **A** : function design, **B** : R calibration, **C** : Calculation from data

| Functions | Biological meaning | Unit | Reference |
| --- | --- | --- | --- |
| $r(x, CO_2)$ | Phytoplankton growth rate | $\text{day}^{-1}$ | A |
| $\pi_p(CO_2)$ | Carbon fixation rate by photosynthesis | $\text{pg C.cell}^{-1}.\text{day}^{-1}$ | a |
| $S_r(x)$ | Sensitivity of phytoplankton mortality to calcification | $\text{pg C.cell}^{-1}.\text{day}^{-1}.\mu\text{mol.L}^{-1}$ | A b e |
| $c(x, CO_2)$ | Calcification rate | $\text{pg C.cell}^{-1}.\text{day}^{-1}$ | A a b c d |
| $S_c(x)$ | Sensitivity of an x-calcification to acidity | $\text{L}.\mu\text{mol}^{-1}$ | A a |
| $\beta(CO_2)$ | Carbon partitioning | $\mu\text{mol.L}^{-1}$ | B b |
| $a(c(x, CO_2))$ | Attack rate of zooplankton on phytoplankton | $\text{Z}^{-1}.\text{mL}.\text{day}^{-1}$ | C f |
| $e(c(x, CO_2))$ | Phytoplankton to zooplankton conversion efficiency | | C g |
| $m(CO_2)$ | Mortality rate of zooplankton | $\text{day}^{-1}$ | C h |

**Table 2.** Model parameters **a** : Sett et al., 2014, **b** : Bach et al., 2015, **c** :Bach et al., 2012, **d** : Bach et al., 2011, **e** : Quéré et al., 2005, **f** :Harvey et al., 2015, **g** : Haunost et al., 2021, **h** : Cripps et al., 2014, **i** : Strom et al., 2018; **A** : function design, **B** : R calibration, **C** : Calculation from data

| Parameters | Biological meaning | Unit | Value / Reference |
| --- | --- | --- | --- |
| $\mu_1$ | Maximum rate of carbon fixation | pg C.cell<sup>-1</sup>.day<sup>-1</sup> | 7.5 / a |
| $K_1$ | Photosynthesis half-saturation constant | $\mu\text{mol.L}^{-1}$ | 1.8 / a |
| $\mu_2$ | Conversion constant of [CO<sub>2</sub>] to [HCO<sub>3</sub><sup>-</sup>] | $\mu\text{mol.L}^{-1}$ | 2250 / B c d |
| $K_2$ | Conversion constant of [CO<sub>2</sub>] to [HCO<sub>3</sub><sup>-</sup>] | $\mu\text{mol.L}^{-1}$ | 3 / B c d |
| $c_1$ | Availability of C allocated to calcification (constant) | | 0.956 / b |
| $c_2$ | Availability of C allocated to calcification (half-saturation) | $\mu\text{mol.L}^{-1}$ | 704 / b |
| $c_3$ | Conversion constant [CO<sub>2</sub>] to [H<sup>+</sup>] | | 4.15E-4 / B c d |
| $f_1$ | Conversion constant for photosynthesis per day | cell.pg C<sup>-1</sup> | 0.1 / C f i |
| $f_2$ | Calibration constant for calcification function | | 1.92 / C |
| $a_{e1}$ | Attack rate sensitivity to calcification (exponential) | cell.pg C<sup>-1</sup> | 1.90E-1 / C f |
| $a_{e0}$ | | Z<sup>-1</sup>.day<sup>-1</sup>.mL | 2.97E-1 / C f |
| $a_{s1}$ | Attack rate sensitivity to calcification (sigmoid) | | 46.35 / C f |
| $a_{s0}$ | | Z<sup>-1</sup>.day<sup>-1</sup>.mL | 14.51 / C f |
| $e_{e1}$ | Conversion sensitivity to calcification (exponential) | | 4.63E-1 / C g |
| $e_{e0}$ | | | 3.13E-2 / C g |
| $e_{s1}$ | Conversion sensitivity to calcification (sigmoid) | | 8.65 / C g |
| $e_{s0}$ | | | 3.23E-1 / C g |
| $s_{r1}$ | Directing coefficient of S<sub>r</sub>(x) | | 0.018 / A a |
| $s_{r2}$ | Ordinate at origin of S<sub>r</sub>(x) | pg C.cell<sup>-1</sup>.day<sup>-1</sup>$\mu\text{mol.L}^{-1}$ | 1.5E-2 / A a |
| $s_{c1}$ | Amplification factor of S<sub>c</sub>(x) | L.$\mu\text{mol}^{-1}$ | 60.37 / A b |
| $s_{c2}$ | Calibration on x | | 0.5 / |
| $s_0$ | Ordinate at origin of phytoplankton mortality | pg C.cell<sup>-1</sup>.day<sup>-1</sup> | 0.1 / |
| $m_1$ | Directing coefficient of m(CO<sub>2</sub>) | L.$\mu\text{mol}^{-1}$.day<sup>-1</sup> | 1.86E-3 / C h |
| $m_0$ | Ordinate at origin of m(CO<sub>2</sub>) | day<sup>-1</sup> | 3.26E-3 / C h |
| $k$ | Competition factor | P<sup>-1</sup>. day<sup>-1</sup> | 3.96E-4 / C |

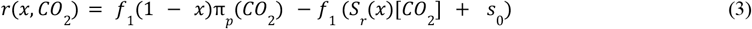

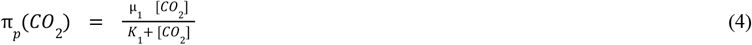

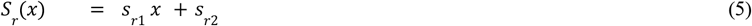

With *r(x, CO*_*2*_) the intrinsic growth rate of coccolithophorids (eq. 3), **π**_*p*_*(CO*_*2*_) the energy from photosynthesis (eq. 4) and *S*_*r*_*(x)* the sensitivity of growth to acidification (eq. 5). Note that when CO_2_ = 0, *r(x, 0)* equals -*f*_1_ *s*_0_ < 0 so that phytoplankton populations decline when photosynthesis is limited. Non-calcifying phenotypes suffer from *f*_1_ (*s*_*r*2_ [*CO*_2_ ] + *s*_0_ ) mortality. The model qualitatively captures variations of intrinsic growth rates obtained from the data (Figure 2a) though the CO_2_ niche generated is more restricted than observed (Sett et al. 2014). Nevertheless, at fixed *x*, the growth rate rises up to a maximum obtained for a CO_2_ concentration almost identical to that of the maximum *r* observed by Sett et al. Past this value, higher CO_2_ concentrations lead to higher phytoplankton mortality and to lower intrinsic growth rates. At fixed CO_2_ of 15 µmol.L^-1^ (the current average value in the global ocean), the intrinsic growth rate decreases as the energy investment in calcification increases. In other words, higher *x* in these conditions leads to less energy invested in growth and to a greater sensitivity to acidity.

**Figure 1.**
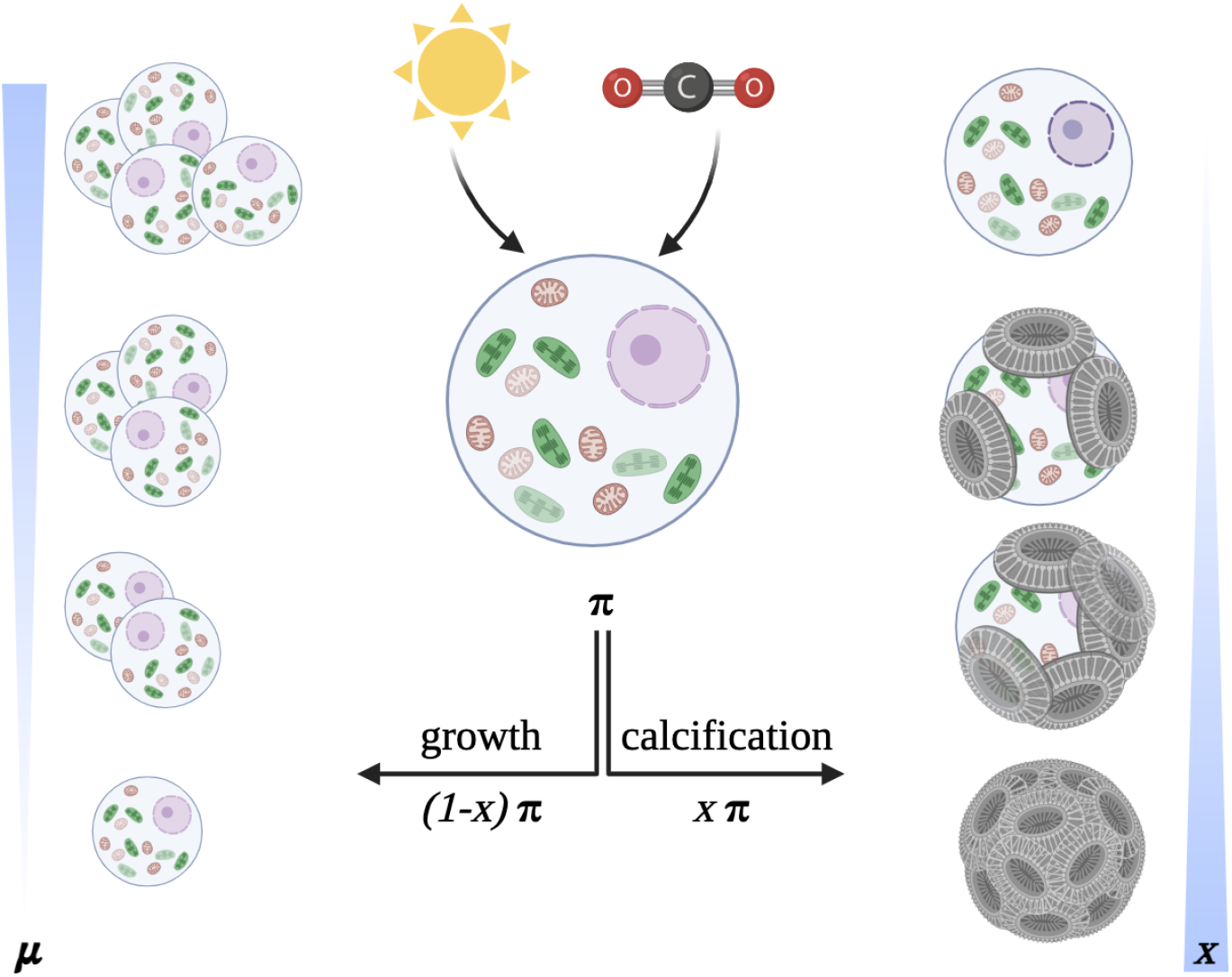
Conceptual principle of partitioning energy from photosynthesis between growth and calcification, with *x* the value of the phenotypic trait characterizing this partitioning, ***μ*** growth and **π** photosynthesis.

**Figure 2.**
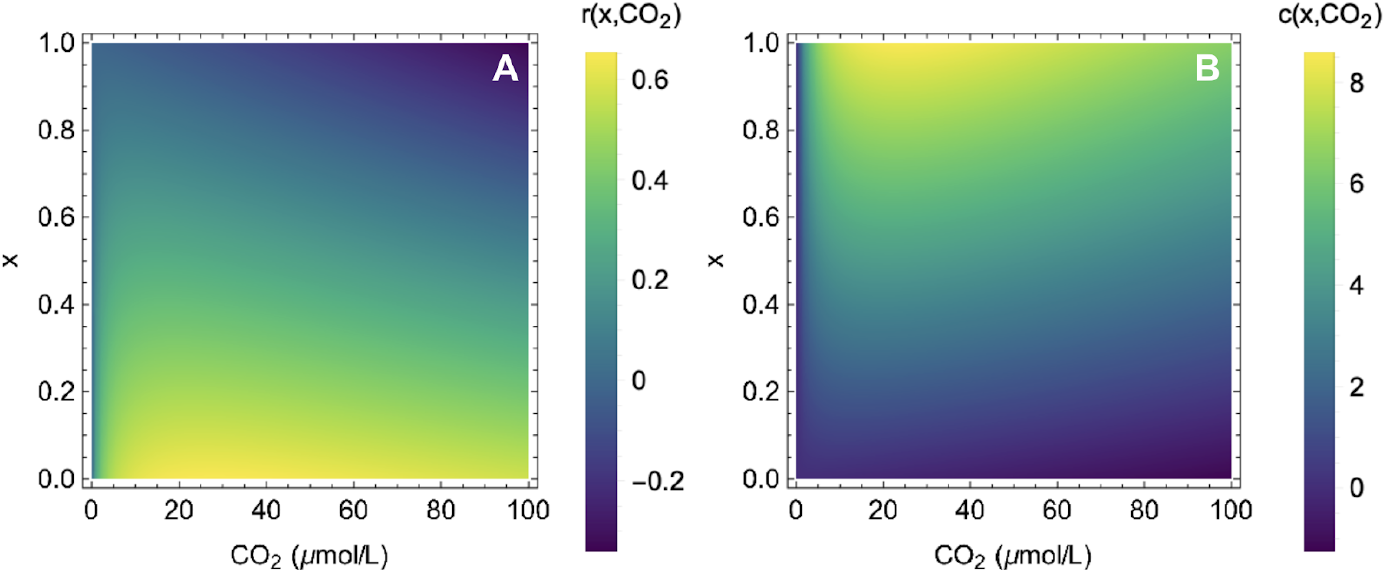
**(A)** Intrinsic growth rate (per day) and **(B)** calcification rate (in pg of carbon per cell per day) of coccolithophorids investing *x* in calcification under different acidity conditions.

#### 1.2.2 The calcification rate

The calcification rate function of a cell (in pg of carbon fixed as calcite per unit time, here per day) is subsequently defined by *c(x, CO*_*2*_) (eq. 6), increasing with the investment *x* in calcification (Table 2 ). This function is adapted from Gafar et al., 2018 and optimized taking into account the variability of trait *x* on data from Sett et al., 2014:

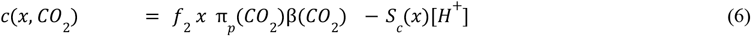

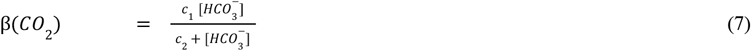

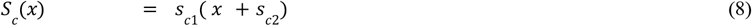

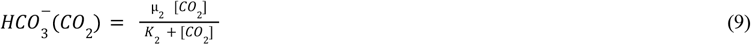

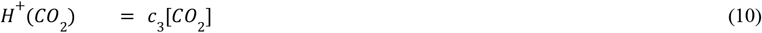

Calcification rate (eq. 6) increases with *x*, π_*p*_, and with the concentration β of carbonaceous substrates in the vicinity of the coccolith-synthesizing golgi vesicles (eq. 7), while acidity leads to a cost for calcification that depends on sensitivity *S*_*c*_*(x)* of calcification to pH, an increasing function of *x* (eq. 8). A high-calcifier phenotype has therefore less resources left to counteract acidity effects on coccoliths dissolution. Since *c(x, CO*_*2*_) is a rate, calcifiers can suffer from net dissolution (negative calcification) once the balance between available energy and pH is unfavorable.

The proposed model correctly reproduces (in value and trend) (**Figure 2**) the measurements obtained on *Emiliania huxleyi* Sett et al., 2014, for an energy investment *x* in calcification currently between 40% and 50% of the cell’s total energy (Monteiro et al., 2016).

#### 1.2.3 Attack rate and conversion efficiency of zooplankton

The attack rate *a(c(x, CO*_*2*_)) is taken to be the number of phytoplankton individuals calcifying at *x* consumed per zooplankton individual per time unit. Because coccoliths are supposed to protect from grazing pressure (Harvey et al., 2015; Kolb and Strom, 2013; Monteiro et al., 2016; Young et al., 2009) we assume that intense-calcifiers are less predated than phenotypes which do not secrete many coccoliths. In addition, zooplankton acquire energy from prey digestion and therefore suffer from energy investment to dissolve carbonate tests (Haunost et al., 2021). To reflect these physiological effects, the conversion factor *e(c(x, CO*_*2*_)) which translates the number of zooplankton individuals produced for the consumption of x-calcified prey is assumed to decrease with the protection of prey. Because trade-off shapes largely affects eco-evolutionary outcomes (De Mazancourt and Dieckmann, 2004), we study the effects of varying trade-off shape by considering two flexible functions (eq.11-14), using data from Harvey et al., 2015; Haunost et al., 2021) where *a* or *e* have decreasing exponential or decreasing sigmoïdes shapes with phytoplankton calcification at *x*, which leads to four scenarios :

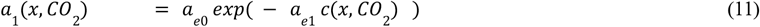

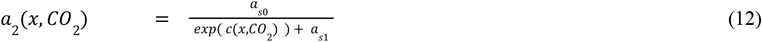

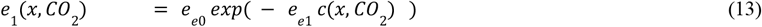

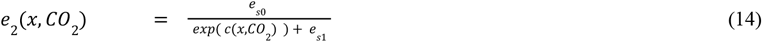

Figure 3 shows that, at constant CO_2_, the attack rate and the conversion efficiency decrease with the rate of phytoplankton calcification (therefore with *x*), consistent with our initial hypothesis. For a given investment *x* in calcification, *a* and *e* initially decrease and then increase with increasing partial pressures. At low CO_2_ concentrations, phytoplankton calcification rates increase, which has a negative effect on *a* and *e*. Beyond this threshold, calcification suffers from increased OA, allowing higher zooplankton grazing and conversion rates.

**Figure 3.**
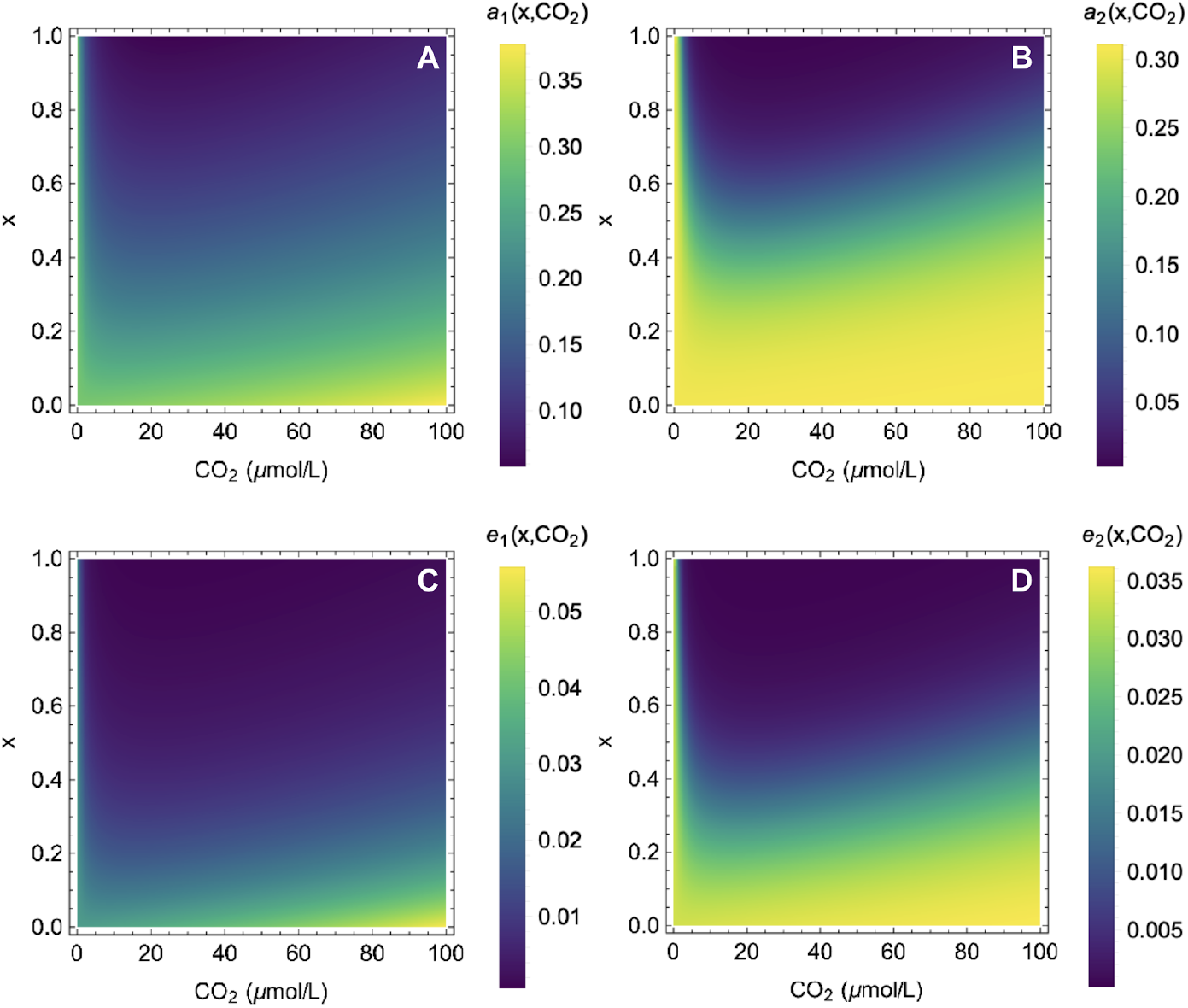
Effects of prey calcification and acidification on zooplankton attack rates and conversion efficiencies.

#### 1.2.4 Zooplankton mortality rate

The zooplankton mortality rate is modeled using an increasing linear *m* function (data from Cripps et al. 2014):

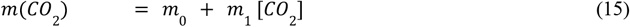

High CO_2_ here increases mortality rates, reflecting osmotic and electrochemical effects of acidification on metabolism.

### 1.3 The eco-evolutionary model

#### 1.3.1 Fitness definition according to adaptive dynamics

Evolution of *x* is studied using adaptive dynamics (Dieckmann and Law, 1996; Geritz et al., 1998; Metz, 1992). Assuming a resident population *P* of trait *x* at the ecological equilibrium, we assume that mutants of trait *x*_*m*_ and population *P*_*m*_ mutants with low amplitude mutations (*x*_*m*_ close to *x*) may appear by chance in the resident population. The main hypothesis of adaptive dynamics is that ecological processes are faster than evolutionary processes. In other words, the ecological equilibrium of the system is reached before the next mutation occurs. *P*_*m*_ is supposed small, as mutant phytoplankton are rare, so that the mutant population develops in an environment constrained by the resident equilibrium *P\**. These mutants can invade the resident population if their growth rate in these conditions is positive (invasion fitness, Metz, 1992). This defines their relative fitness *ω(x*_*m*_, *x)* (eq.16) :

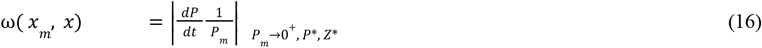

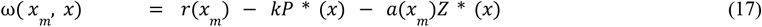

Evolution of phenotypes *x* can then be approximated by the canonical adaptive dynamics equation (Dieckmann and Law, 1996) :

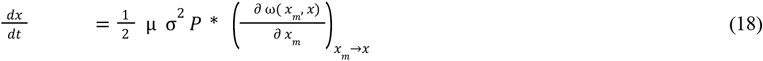

Where µ is the per individual mutation rate, σ the amplitude of the mutation process and 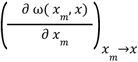 the fitness gradient constraining the direction of evolution. Evolution stops once *x*^*^the evolutionary singularity is reached 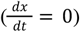, *i.e*., when the population goes extinct or the fitness gradient cancels.

#### 1.3.3 Convergence and invasibility criteria

Dynamics close to the evolutionary singularities are determined based on convergence and invasibility criteria (Dieckmann and Law, 1998; Geritz et al., 1998). Convergence properties assess whether, in the vicinity of singular strategies, phenotypes closer to the singularity have an advantage.

Evolutionary dynamics converge to x* when : 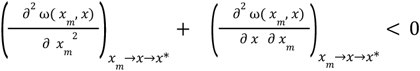

Strategy *x\** cannot be invaded (non-invasibility criterion, ESS in Smith and Price 1973 by nearby mutant strategies when :

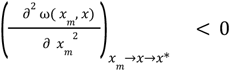

When the singular strategy is convergent and non-invasible, it is a continuously stable strategy (CSS in Eshel, 1983). In such situations, phytoplankton populations eventually invest *x*\* in their calcification and cannot be invaded by populations with a different *x* phenotype. Conversely, when the singular strategy is non-invasible but non-convergent, it is called a Repellor and the evolution of the trait then moves it away from *x*\*. Phenotypes starting at low calcification levels then lead to a complete counterselection of calcification, while phenotypes starting at high calcification will lead to evolutionary increase in calcification.

### 1.4 Simulations

#### OA scenarios

Ocean acidification is strongly correlated with the increase in atmospheric CO_2_ concentration. Nonetheless, local variations in seawater CO_2_ may largely vary in amplitude and in time (Orr et al., 2022, McNeil and Matear, 2008). Taking this into consideration greatly complexifies the present study without giving a more realistic overview of the eco-evolutionary impacts of ongoing OA. Therefore, we consider homogeneous seawater CO_2_ concentrations and carry out rising OA simulations according to the present RCP scenarios (Riahi et al., 2011). The most optimistic scenario predicts a fall in CO_2_ levels, stabilizing at 380 ppm in 2100 (RCP2.6). The increasing scenarios differ in their intensity. CO_2_ would rise from 510 ppm in 2100 to 530 ppm in 2200 in the RCP4.5 scenario, while in 2100 it would be around 680 ppm in the RCP6 scenario and 1200 in the worst-case scenario RCP8.5. While concentrations would reach a plateau in the 2150s at 750 ppm for RCP6, they would continue to rise until 1950 ppm in 2230 in the case of RCP8.5. The scenarios then represent increasingly strong acidification rates and intensities: a decrease in OA in the RCP2.6 case, and an increase in the intensity of OA between the RCP4.5, RCP6 and RCP8.5 scenarios.

#### Stochastic simulations of evolutionary dynamics

We approximate the variations of CO_2_ concentrations in the surface ocean by instantaneous equilibrium with the atmosphere. For RPC4.5, RCP6 and RCP8.5 scenarios, we estimate the CO_2_ concentration from data using Self-Starting NLs Logistic models. Polynomial approximation is used for RCP2.6. The characterization of eco-evolutionary dynamics using the canonical equation (eqs. 16 to 18) relies on the hypothesis of small and rare mutations. However, experimental evidence suggests that evolution in response to acidification can be quite fast. Fast evolution may help the persistence of a given species (Gomulkiewicz & Holt 1995), but whether this extends to more complex systems remains unclear (Loeuille 2019). To tackle this question, we use stochastic simulations of eco-evolutionary dynamics, where we can explicitly vary the speed of the mutation process. These simulations are based on tau-leaping techniques (Gillepsie et al, 2001). Every **τ** time step, ecological interactions are derived from poisson distributions and on average 5% of mutants arose from new individuals. Mutant traits are drawn from normal distributions (5% of variance), centered around the parent’s trait. 50 simulations to the eco-evolutionary equilibrium under constant OA are carried out to obtain a typical distribution of *x* traits in coccolithophorids populations before OA. The potential carbon loss is estimated by normalizing the potential carbon content in coccolithophorids population by its value when OA starts.

The ecological model calibration and tau-leap simulations were carried out using R software. Symbolic mathematical analyses and graphical outputs were obtained using Mathematica.

## 2. Results

### 2.1 Ecological equilibrium states (no evolution)

At steady state, the system (eq. 1, 2) has three distinct equilibrium points (eq. 19, 20, 21).

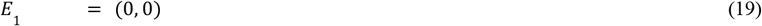

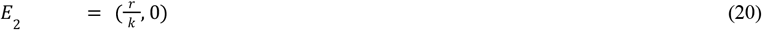

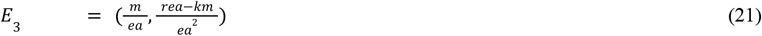

E_1_ is the trivial equilibrium (eq. 19) where neither phytoplankton nor zooplankton survive. This point is stable when the growth rate of phytoplankton is negative.

### 2.2 Phytoplankton dynamics penalized by OA and calcification in absence of herbivory pressure

When *r* is positive, phytoplankton exclude their predators if zooplankton mortality exceeds the total amount of resources extractable from *x*-calcifying haptophytes 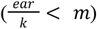 which leads to E_2_ (eq. 20, shown Figure 4, within red dashed limits). Note that in this equilibrium, calcifying phytoplanktons have lower densities *r*/*k* than less-calcifying ones (as calcification reduces *r* (Figure 2)) regardless of the trade-off shape used. Effects of OA follow its consequences on *r* (section 1.2.1 & Figure 2), *i.e*. densities are first increased (higher photosynthesis) then decreased (metabolic costs).

**Figure 4.**
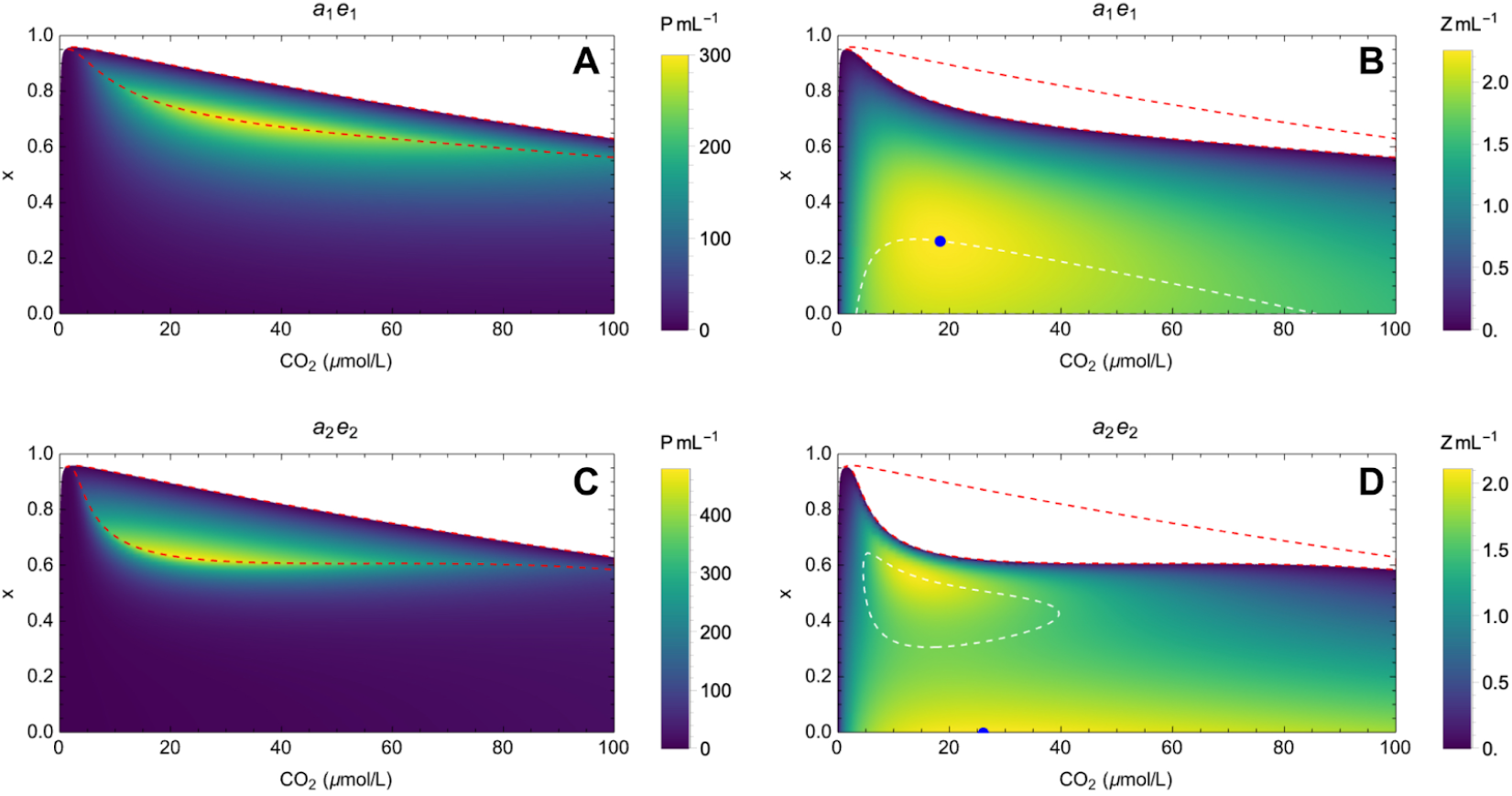
Phytoplankton (A,C) and zooplankton (B,D) density at equilibrium for 2 combinations of trade-off between zooplankton attack rate and conversion factor. White area : no species survive in the environment. Red-outlined area : E_2_ relative space (*x, CO*_2_ ). White-outlined area : *(x, CO*_2_ ) space of E3 where 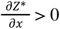. Blue point : maximum zooplankton density. X-axis: CO_2_ concentration in µmol.L^-1^, Y-axis: value of line *x*.

### 2.3 Grazers characteristics control calcification effects on phytoplankton abundance

The coexistence equilibrium E_3_ is stable when it is feasible (eq. 21), which allows grazers to coexist consuming phytoplankton. Partial derivation on *x* of (eq. 21) causes:

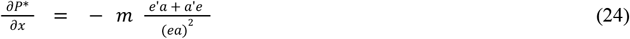

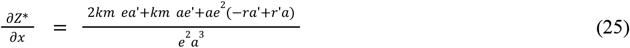

With 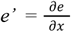 and 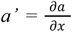. Calcification outcomes on P* densities only depend on how *x* affects zooplankton characteristics. Since *a’* < 0 and *e’* < 0, increased calcification leads to a relaxation of top-down control. Phytoplankton density then increases (eq. 24, Figure 4 left panels) with protection while the impact of calcification on Z* densities seem less obvious to predict (eq. 25). Therefore, calcification benefits phytoplankton only in coexistence situations. Zooplankton generally suffer from an increase in algae protection except in white areas (Figure 5, right panels) where they benefit indirectly: calcification then favors energy transfers within the community. According to E_3_, Z* depends on grazing pressure intensity and on P* 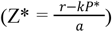. If calcification favors P* densities as shown previously, it nevertheless represents a constraint on grazing pressure. Thus, according to (eq. 21) we understand that an increased calcification involves decreased Z* densities as long as *(-ra’ + ar’)* < 0 since *a’* < 0, *e’* < 0 and 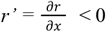 (eq. 25). However, when this ratio between the cost of calcification on intrinsic growth (*r’*) and its cost on attack rates (*a’*) becomes sufficiently positive, 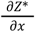 can then become positive as well. When grazing pressure suffers more than algal growth to the increase of their protection, then Z* density increases at equilibrium and zooplankton populations benefit from calcification. These non-trivial effects come from an increase of P* densities with an attack rate decrease, which is ultimately favorable to zooplankton. Under exponential tradeoff shapes (Figure 4, upper panels, white dashed areas) non-trivial effects are observed in wide conditions and interestingly zooplankton prefer calcified algae (max densities at x ≈ 0.3). Under sigmoïde tradeoff (lower panels) non-trivial effects are more constrained but fits best with observed phenotypic diversity and actual environmental conditions. From the outset, acidification seems to have more subtle ecological effects since it acts on *m, r, e* and *a* in a non-monotonic way (see Results, 2.4)

**Figure 5.**
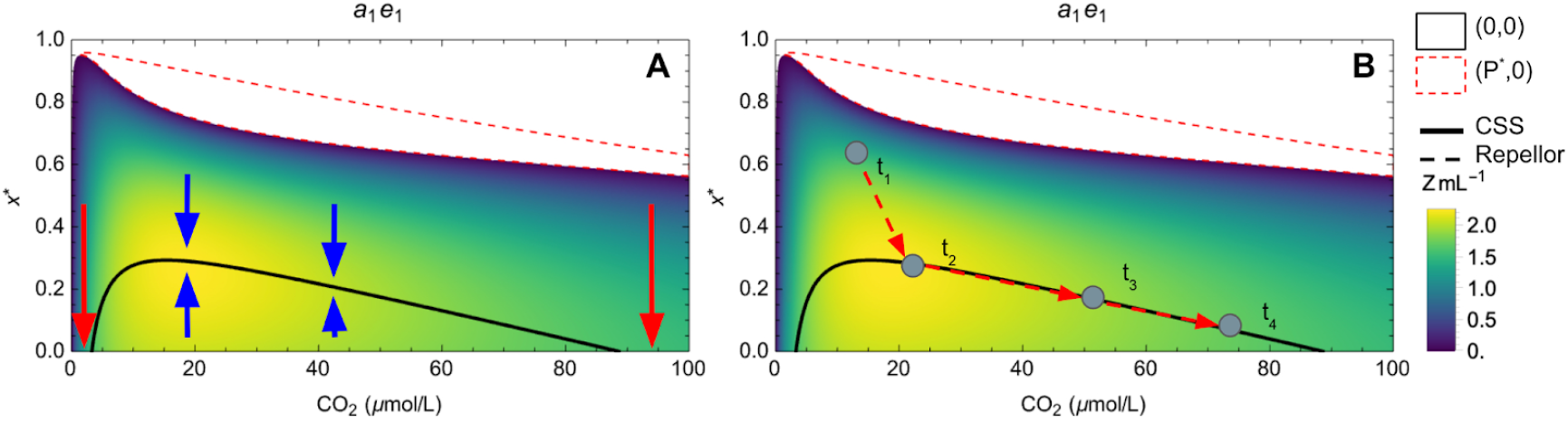
E_3_-diagram representing the selected strategy (solid line, CSS) in the case of a decreasing exponential attack rate in an acidifying environment. The conversion efficiency e1 is exponentially decreasing. Arrows : direction of change. A fictive scenario of the evolution of the value of *x* over time (B) is shown for ease of understanding. Zooplankton densities are also shown. Red dashed outlined area : (*x, CO*_*2*_) space excluding zooplankton. White area : (*x, CO*_*2*_) space where no species is able to survive. All other parameters are fixed at the values shown in Table 2.

### 2.4 Ecological impacts of OA

Analytical resolution shows that sensitivity of attack rates and conversion efficiency to an increase in CO_2_ is the key parameter which determines phytoplankton densities variations. From (eq. 21) we obtain that:

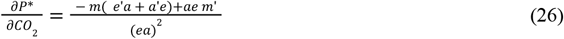

For low CO values, 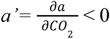 and 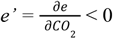 as long as the calcification rate increases with carbon availability (Figure 3). When calcification suffers from OA then *a’*> 0 and *e’* > 0 (Figure 3, high CO_2_ values). Since *a, e* and *m* are positive and *m’* > 0, when *a’* < 0, *e’* < 0 (in relatively high seawater pH) then 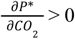. This means that OA will first benefit phytoplankton populations under coexistence due to the fertilizing effect of CO_2_ (higher primary production) but once OA costs on physiology exceed its fertilizing effect, then a’ and e’ become positive and 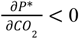.

There is an optimum of intermediate CO_2_ for which energy flows within the system and zooplankton densities are maximum (Figure 4, right panels) corresponding to lower seawater acidity than today. When zooplankton is able to survive Z* benefits from fertilizing effects of OA on P* before decreasing due to toxicity. From (21), we obtain that OA induces:

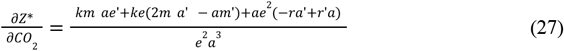

Up to an intermediate CO_2_ concentration, OA has negative effects on attack rates, conversion factor and positive effects on phytoplankton growth as described before (*a’* < 0, *e’* < 0 and *r’* > 0) and the opposite effect for higher concentrations. In the fertilization area, OA increases grazers densities if its benefit on phytoplanktonic growth rate is much greater than its cost on the attack rate (*ra*′ ≪ *r*′*a*). In the toxicity area, calcification suffers from OA, which favors the attack rate of zooplankton on phytoplankton and therefore reversed dynamics. These results are consistent with the initial physiological hypotheses, namely that intrinsic phytoplankton growth and calcification follow a CO_2_-structured niche.

### 2.5 Evolution of calcifying capacities under current oceanic conditions

Under current ocean acidity, phytoplankton are evolving towards calcifying or non-calcifying abilities (Figure S1), depending on initial phenotypic conditions (Figure S2). Evolutionary outcomes slightly differ between exponential (low calcifiers selected in Figure S1 top, S2 left) or sigmoïde (high or non-calcifiers selected in Figure S1 bottom, S2 right) trade-off for attack rates. These results can be explained by considering the fitness gradient of a mutant *x*_*m*_ relative to a resident *x* (eq. 17):

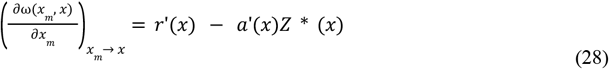

Calcifying singular strategies cancel this fitness gradient and can be understood as the balance between the costs of calcification on growth (r’(x) < 0 ) and the benefits due to herbivory pressure reduction (-a’(x)Z*(x) > 0). The evolutionary outcome of the system is then discriminated according to the convergence and invasibility criteria and simulated (Figure S2 for instance). At current CO_2_ levels (15 µmol.L-1), low calcifiers (CSS x* ≈ 0.3) optimize this cost/benefit balance because great protection effects are already present for small x (attack rate is exponentially decreasing) whereas high-calcifying (CSS x* ≈ 0.7) or non-calcifying strategies are selected depending on initial phenotypic conditions when a is sigmoïde (low x : high growth/low protection, high x : small growth/high protection). Interestingly, these results emerge from (i) modal effects of CO_2_ depending on its concentration. At low levels, CO_2_ is mostly invested in growth and reproduction, so that any investment in calcification is counterselected (Figure S1,S2) and at higher levels, acidity counterselects calcification (see Supplementary). (ii) the selective herbivory pressure of zooplankton (see Supplementary). Here herbivory pressures maintain the capacity of phytoplankton to calcify and that the coexistence of populations within trophically structured communities is therefore a determining factor in the oceanic carbon cycle *P\** will be low (see Figure 4, left panels) and so will *Z\** (see Figure 4, right panels) which does not select for calcification.

### 2.6 Evolution of calcification maintains coexistence under OA

To understand how *x* evolves with acidification, we use Ecology-Evolution-Environment diagrams (E3-diagrams, Ferriere and Legendre, 2013), for which we recover the value of the evolutionary singularities in an increasingly CO_2_-rich environment (Figure 5&6). In all cases (i) calcification is counter-selected at low CO_2_ (low availability of energy) and at high CO_2_ (high costs of calcification due to acidification) (red arrows on Figure 5 and 6); (ii) at intermediate CO_2_, a certain investment in calcification (ie, a CSS) is selected (blue arrows Figure 5 and 6). Substantial differences then exist between sigmoid and exponential trade-offs. While exponential attack rates typically lead to low calcification (around 0 to 0.3 on Figure 5), sigmoid shapes lead to very high investments (around 0.6 on Figure 6) that coexist with an alternative “no calcification” strategy. Populations originally investing less than the *x\** given by the dotted curve (repellor) will evolve towards a loss of their ability to calcify (blue arrows).

**Figure 6.**
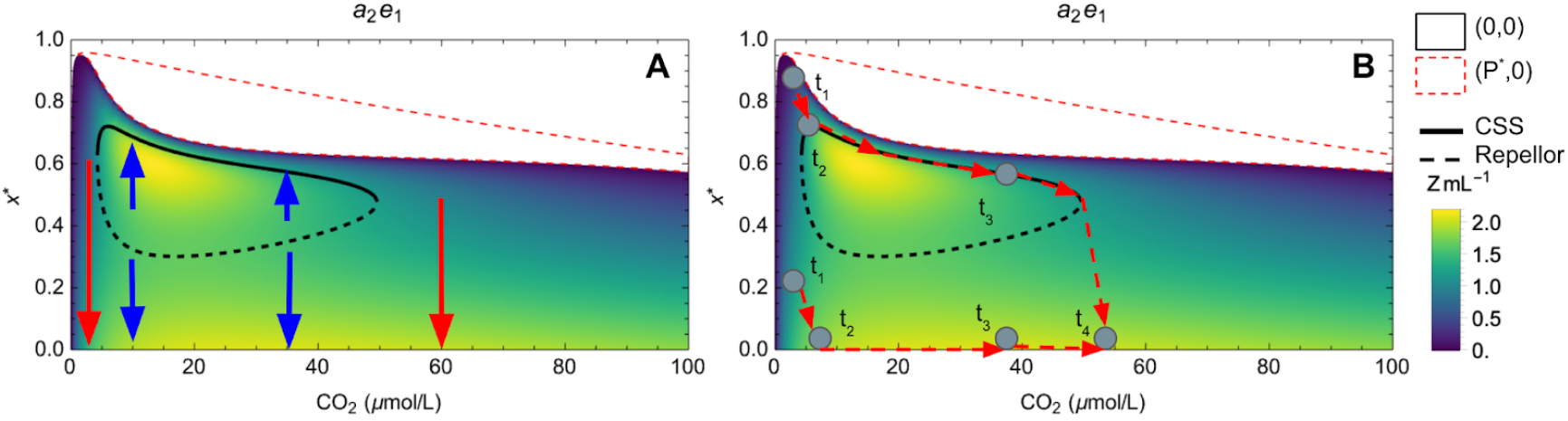
E_3_-diagram : selected strategy (continuous line : CSS, discontinuous line : Repellor) in the case of a decreasing sigmoidal attack rate in an acidifying environment. The conversion factor is exponentially decreasing. Arrows : direction of evolution. A fictive scenario of the evolution of *x* over time (B) is represented for ease of understanding. Zooplankton densities are also represented. Light gray area : (*x, CO*_*2*_) space excluding only zooplankton. Dark gray area : (*x, CO*_*2*_) space where no species is able to survive. All other parameters are fixed at the values shown in Table 2.

Let us now turn to the effects of a progressive increase in CO_2_ (acidification) (broken red paths on figures 5 and 6) and how the resulting eco-evolutionary dynamics change the trophic structure and energy transfers. At low CO_2_ the coccolithophorids lose their ability to calcify which leads to low *P\** and *Z\** densities (Figure 4). At intermediate CO_2_ (from 4 to 20 µmol.L^-1^) evolved calcification increases, which is linked to higher *P\** and *Z\** densities. Photosynthesis is then close to its optimum and sustains high zooplankton densities. This in turn leads to high protective benefits of calcification, therefore to its maintenance. The evolution of calcification here promotes energy transfers. We can also see that these arrows always point away from the exclusion zones, which shows that the evolution of calcification favors coexistence. Further acidification causes a decrease in evolved calcification (t_2_ to t_4_ path on Figure 5), then to its loss (here when CO_2_ reaches 90 µmol.L^-1^). Energy costs on growth *r’(x)* (eq. 28) are too high to maintain coccoliths even though their protective benefit remains substantial (*-a’(x)Z*(x)* strong). When the attack rate is decreasing sigmoidally (Figure 6), qualitative dynamics are consistent with the previous case at low CO_2_. Maximum calcification is evolved at intermediate CO_2_ (15 µmol.L^1^) though populations with initially low investment will lose their ability to calcify (blue arrows). Thus, for concentrations between 5 and 50 µmol.L^-1^, two evolutionary dynamics coexist, calcified and non calcified.

Importantly, this coexistence of two eco-evolutionary outcomes leads to vastly different dynamics under acidification. At its optimum (Figure 6), the system is not very sensitive (evolved calcification changes little, t_2_ to t_3_ on Figure 6) and resilient to acidification, until a threshold acidification value is reached. At around CO_2_ = 50 µmol.L^-1^, the repellor and CSS singularities collide. A critical transition point (evolutionary tipping point) is reached and a very slight acidification leads to a sudden loss of the ability to calcify (t_3_ to t_4_ on Figure 6). This evolution of low calcification leads to high grazing and tends to promote trophic transfers so that *Z\** densities increase. Calcification cannot be easily recovered because eco-evolutionary dynamics are trapped in the repellor basin of attraction (a case of eco-evolutionary hysteresis). The results of other trade-off combinations are qualitatively coherent with the ones we describe here (see supplementary)

#### 3-OA might endanger coccolithophores ability to calcify affecting calciferous carbon input in the ocean

To further investigate the interplay of the three timescales (ecology, evolution and changes in CO2 concentration), we conducted tau-leap simulations. The more the OA is intense (by amplitude and speed), the more it affects the ability to calcify (Figure 8). It can decrease by a few percent over long periods under the RCP4.5 and RCP6 scenarios, while an OA under RCP8.5 can lead to a reduction of almost 40% in the average ability to calcify. This sharp decline is due to a drastic drop in population densities, in phenotypic diversity (Figure S4) and an intrinsic evolution of trait *x*. The oceanic carbon cycle is fed in particular by the sinking of carbonates from calciferous tests. When populations are declining and are less able to calcify, this negatively impacts the carbon pump. Thus, over the long term, low or moderate OA (RCP4.5 and RCP6 respectively) have relatively little effect on coccolithophorid densities and their ability to calcify, leading to small decreases in carbon influx (a few percent for RCP4.5, -15% for RCP6). The RCP8.5 scenario could lead to almost the disappearance of coccolithophorids contribution in the carbonate cycle. At the same time, individual trophic transfers from phytoplankton to zooplankton increase and this result is explained by the reduced capacity of coccolithophorids to protect themselves, increasing herbivory pressure.

## Discussion

In this study, we showed that the ongoing acidification of the oceans had different ecological consequences for calcifying communities, depending on whether or not evolution of marine phytoplankton is possible. These results are based primarily on an understanding of the effects of calcification and acidification on community dynamics. The energetic investment in calcification is a balance between energetic investment in growth functions and costs to the protection of the organism (Monteiro et al., 2016). We first showed that calcification promotes phytoplankton densities in a system where zooplankton coexist. While calcification often reduces zooplankton abundances, there are conditions under which it increases it. Acidification promotes transfers between the two trophic levels up to a certain level. Past this concentration, it penalizes coexistence and trophic interactions. These ecological patterns change when calcification is allowed to evolve. For low CO_2_ concentrations calcification is counter-selected and energy transfers are restricted. Higher CO_2_ eventually selects for calcification that can be either maintained at high levels and relatively insensitive to acidification (decreasing sigmoid modeling) or increase progressively to intermediate levels but be relatively sensitive (exponential modeling). High acidification always leads to the loss of calcification thereby increasing energy transfers from phytoplankton to zooplankton. These results are summarized in Figure 7.

**Figure 7.**
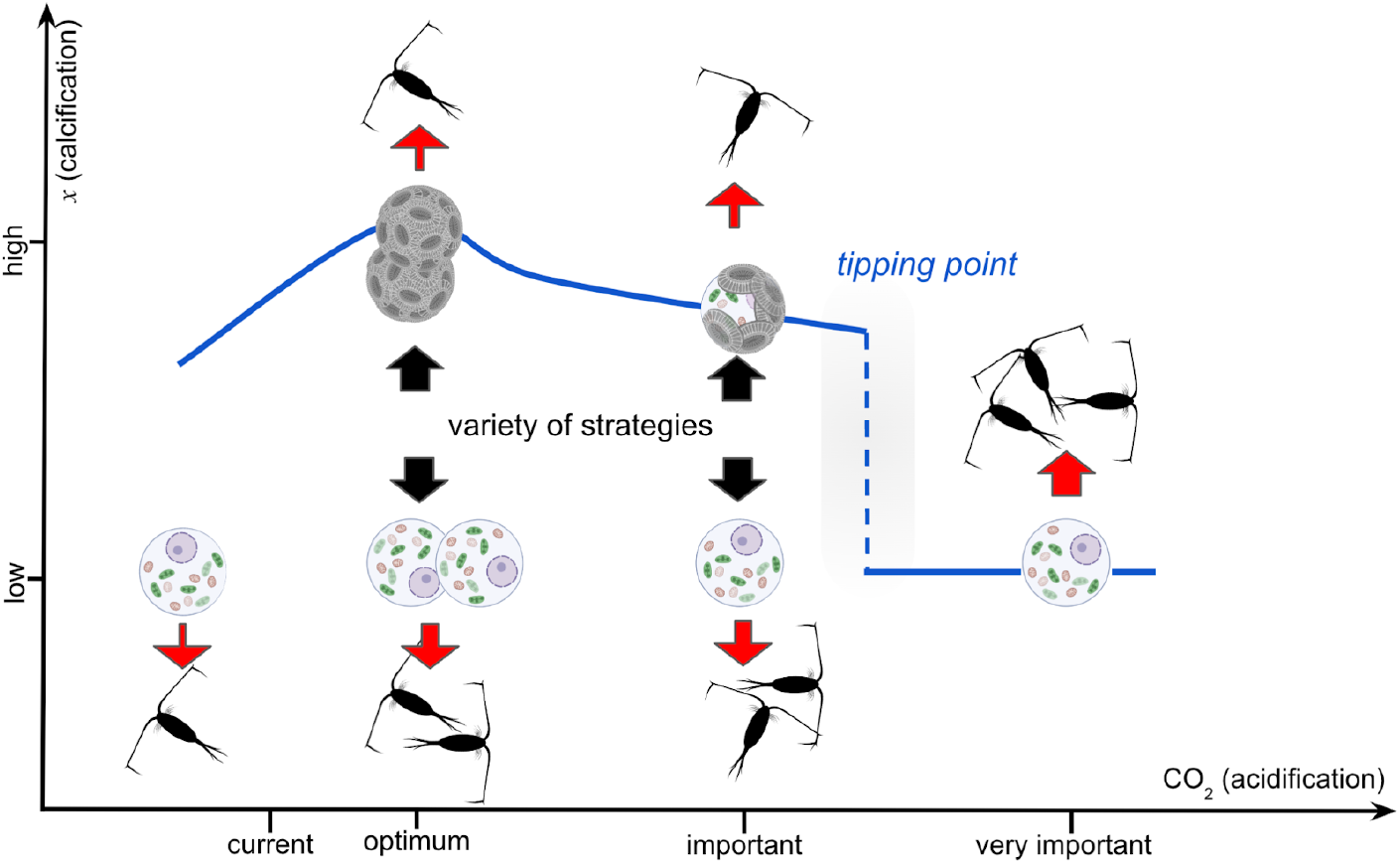
Summary of the effects of ocean acidification on the evolution of calcification and its implications for energy transfer. The study showed a sigmoid relationship for the attack rate seemed the most likely and is prefigured here.

**Figure 8.**
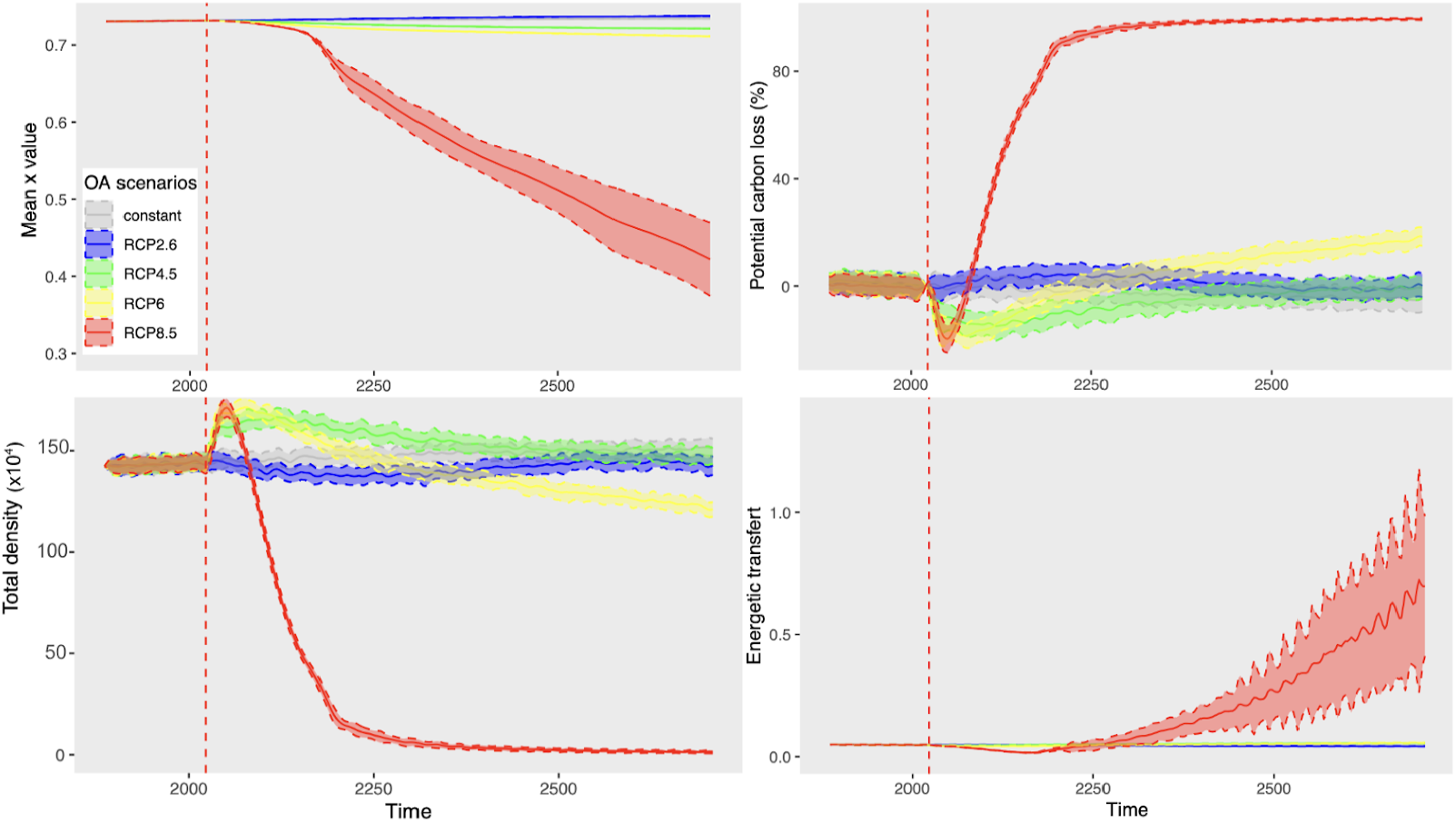
Impacts of RCP scenarios on coccolithophorids eco-evolutive dynamics and implications for the carbon cycle and energetic transfers. Full line : mean of 50 simulations for every RCP scenario. Dashed-line : standard deviation.

We first showed that calcification promotes phytoplankton abundance at equilibrium by alleviating the top-down control of their herbivores. These results can be related to what has been observed for terrestrial communities (Leibold, 1996). Higher calcification then leads to lower abundances of zooplankton. Nevertheless this effect needs to be put into perspective as we have seen that there are conditions for which marine zooplankton develop more when algal calcification increases. These results should be seen in relation to the diversity of effects of calcification on zooplankton dynamics found in the literature (Harvey et al., 2015). In our case, this non-trivial consequence comes from the fact that under certain conditions, calcification penalizes the attack rate of zooplankton more than the intrinsic growth of coccolithophorids. Under these conditions, the density of phytoplankton increases even more, which enhances energy transfers. We observed this phenomenon in a relatively restricted range of *x* with CO_*2*_ concentrations relatively close to current concentrations. In this range, calcification promotes energy transfers and the productivity of marine ecosystems. Excessive calcification is however detrimental to the maintenance of the community as it eventually leads to the exclusion of zooplankton.

Acidification affects phytoplankton and zooplankton populations in a similar fashion. For a relatively low level of acidification (around 20 µmol.L^-1^ of CO_2_), phytoplankton and zooplankton benefit more from the increase in the carbon substrate in the environment, as this increase of energy matches physiological costs due to the lowering of water pH. Since the ocean currently contains an average of 15 µmol.L^-1^ of CO_2_, this phenomenon could be observed until around 2050 in a ‘business as usual’ scenario (Riahi et al., 2011). When the CO_2_ concentration exceeds this threshold, the balance between the benefits of increasing the carbon substrate for trophic transfers in the system and the costs of acidification is reversed. Without changes in coccolithophorids, acidification then reduces trophic transfers within the community. All of these results are qualitatively robust, whatever the form of the trade-off used to model the attack rate and the conversion efficiency.

If evolution of phytoplankton is possible, as suggested by various experimental works (Kroeker et al., 2013; Schlüter et al., 2014), we note that the resulting eco-evolutionary dynamics constrain the resilience and functioning of the system in various ways. We then took into account the evolution of phytoplankton in our system. We modeled the energetic investment x in calcification, using adaptive dynamics methods. Exponential attack rate scenarios led to only one calcifying strategy at evolutionary equilibrium, obtained for *x*\* ≈ 0.3 under current pH conditions. Current observations suggest an average *x* ≈ 40% - 50% (Monteiro et al., 2016). Our sigmoid scenarios typically showed selected values that were a bit higher, though the exact positions of the evolutionary singularities in both scenarios likely depend on the parameters we estimated to build the model. Sigmoid trade-offs also lead to the coexistence of contrasted evolutionary outcomes (high vs low calcification). These results are more in line with observations on the diversity of calcification strategies currently observed (Monteiro et al., 2016), with some species investing heavily in calcification and others with little or no protection.

Ocean acidification directly affects the physiological capacity to calcify (Sett et al., 2014), leading to a reduction in calcification over generations in coccolithophorids (e.g. (Schlüter et al., 2014). Kroeker et al., 2013 showed that acidification can reduce the ability to calcify by up to 23% in coccolithophorids by 2100. Our study is in line with these results, since whatever the form of the trade-off, acidification causes a reduction of evolved investment in calcification. When the attack rate is modeled by a decreasing sigmoid relationship, this counterselection occurs abruptly (eco-evolutionary tipping point) for CO_2_ concentrations of around 50 µmol.L^-1^. Such an abrupt shift here completely changes trophic structures, largely increasing the trophic flux thereby likely affecting the energetic basis of marine systems. Following global scenarios, these concentrations could be reached by 2150 in the worst-case scenario (Riahi et al., 2011). Such a tipping point would also lead to hysteresis: past the loss of calcification, coccolithophorids are not expected to recover their ability to calcify when pH rises again. This result is quite original because it is an evolutionary tipping point that rubs off with the classical description of ecological tipping points (Dakos et al., 2019; Scheffer and Carpenter, 2003).

Interestingly, consequences of acidification on the structure of phytoplankton-zooplankton communities depend on whether evolution of the ability to calcify is possible or not. In evolutionary scenarios, the loss of calcification due to acidification tends to lead to high trophic fluxes and large zooplankton abundances, while the opposite is observed if evolution is too slow (ecological dynamics analysis). This effect shows that evolution maintains the community and ultimately increases energy transfers within the system in response to ocean acidification.

Our results show the importance of taking evolution into account in the development of models studying the response of certain functional groups to climate change. The reduction of coccolithophorid calcification due to ocean acidification is a major issue in the regulation of the carbon cycle. While the ocean absorbs 30% of the CO_2_ released by human combustion (Millero, 1995), coccolithophorids are responsible for transferring carbon to the deep ocean (counter-pumping). This is because their heavy, calcified structures lead to high sinking rates of their CO_2_. By altering calcification, ocean acidification may largely reduce this biological counter-pump thereby enhancing the effects of global warming (Riahi et al., 2011). Nevertheless, our findings on the trophic consequences of the system could balance this effect. Quantitative studies are therefore needed to better understand which of these two effects will likely be dominant in the future CO_2_ cycles.

Despite the relative simplicity of the ecological model (classic Lotka-Volterra), the considerable effort put into determining the physiological responses of these organisms to acidification gives the study a certain robustness. However, one major question remains unanswered. The literature shows that over geological time, several radiative phenomena in the evolution of coccolithophorids have been observed (Monteiro et al., 2016), whereas we do not obtain evolutionary branching conditions in this study. Arguably, the competition factor *k* should have been *x*-dependent as calcification can affect the buoyancy of the cell (Monteiro et al., 2016) which determines its relative position in the water column. Since there is a vertical gradient of light energy opposite to that of CO_2_ concentrations, there is then different competition depending on position and therefore depending on *x*. Two similar phenotypes *x*_1_, *x*_2_ are therefore more in competition than different phenotypes. Were we to include such competitive effects, we expect that limiting similarity would favor the emergence of contrasted strategies (Doebeli and Dieckmann, 2000.; Macarthur and Levins, 1967). Finally, our deterministic approach ignored the stochasticity inherent in these evolutionary systems. We assumed that the speed of evolution was much greater than that of human disturbance. This assumption needs to be put into perspective, and it would have been necessary to look at the outcome of the same stochastic system when subjected to different rates of acidification and evolution.

## Supplementary materials

### Equilibrium feasibility and stability

The Lotka-Volterra dynamic system (eq 1, eq 2) has three distinct equilibriums.

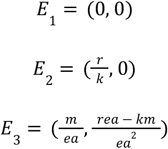

We note 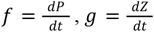 and *J* the associated Jacobian matrix of the system :

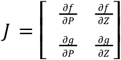

and 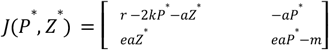

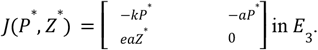

*E*_3_ exists when *Z*^*^ > 0 and *P*^*^ > 0. *P*^*^ > 0 in all cases because *e* > 0, *a* > 0 and *m* > 0.

Therefore, *E*_3_ exists when 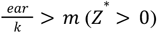. In these conditions,

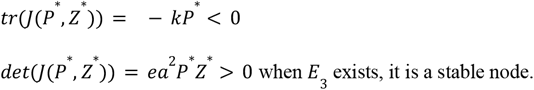

*E*_2_ exists when *Z*^*^ = 0 and *P* ^*^> 0 meaning that *r* > 0 and 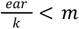. In these conditions,

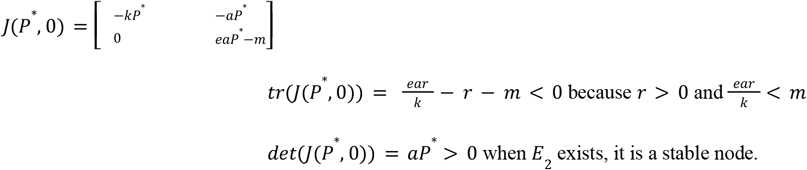

If 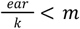 and 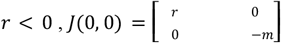

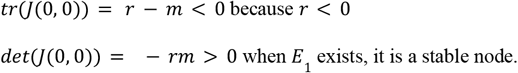

#### Without grazers, calcification and acidification penalize phytoplankton growth

We first consider environmental conditions in which only the phytoplankton is able to persist (E_2_ eq 20). We obtain from Figure 4 (A,B,C,D, red outlined area) that calcification (x-labeled axis) and acidification (CO_2_-labeled axis) lead to a decrease in P* densities regardless of the form of the trade-off shape used. In absence of selective pressure due to herbivory, the reproduction and maintenance of algae suffers from energy losses associated with their protection. After partial derivation on *x* of (eq 20):

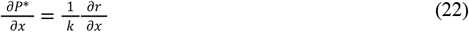

With 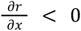 the cost of calcification on intrinsic growth. When grazers are excluded, we confirm that phytoplankton density at equilibrium decrease with their investment in protection. According to (eq 20) acidification causes:

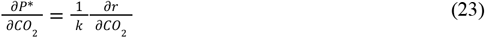

Since 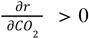 for small CO_2_ values and becomes negative thereafter, the equilibrium density of phytoplankton is expected to vary as its intrinsic growth rate with OA (Figure 2). This analytical result contrasts at first sight with our observations in Figure 4 but remains consistent because there is in fact a small space (*x*, CO_2_) within the exclusion zone for which OA leads to an increase in density. OA penalizes phytoplankton dynamics by affecting the energy turnover associated with its growth and the conservation of high calcification rates to counterbalance coccoliths dissolution.

---

**Figure S1.**
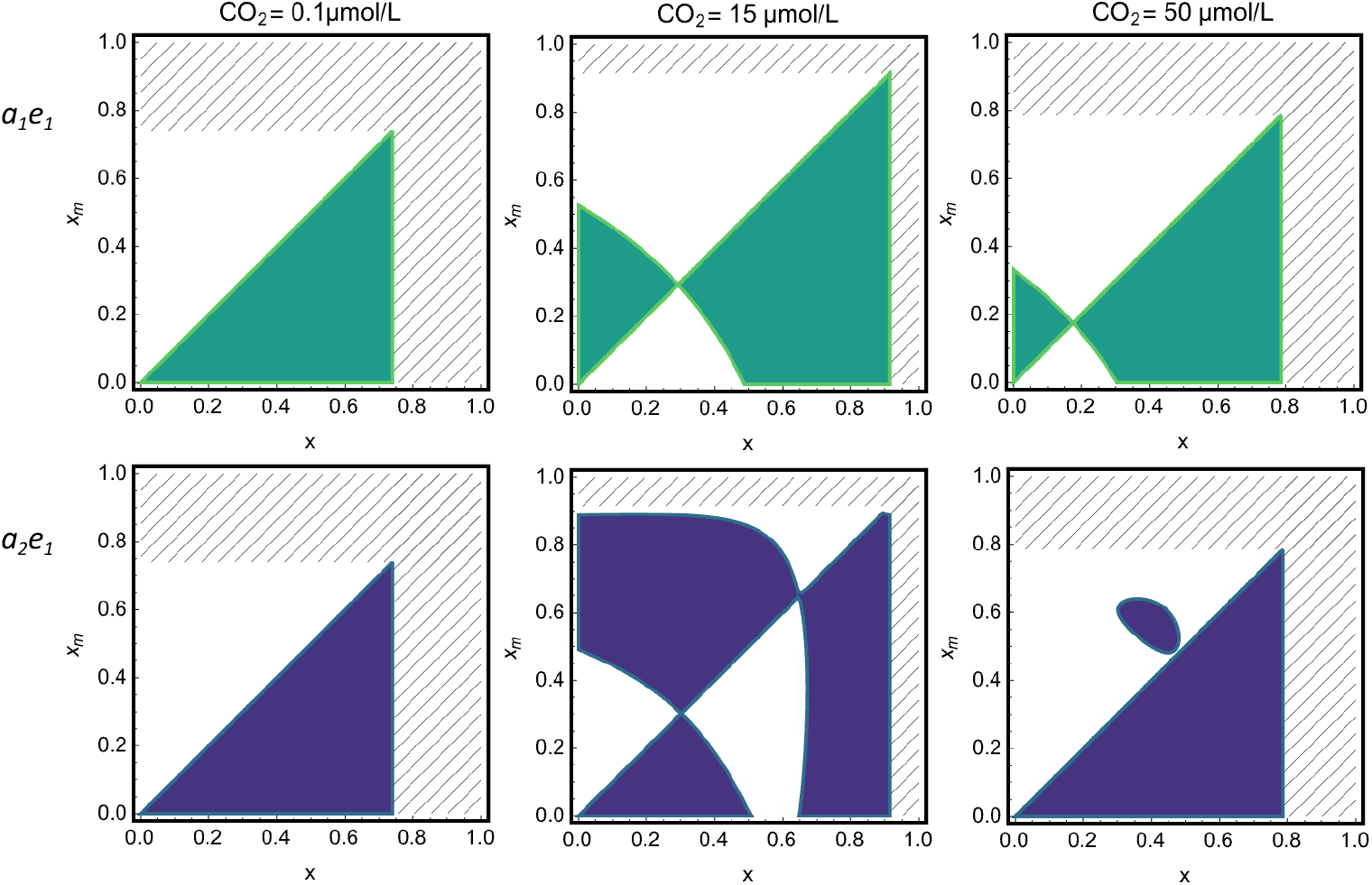
Pairwise Invasibility Plots (PIP) representing the sign of the fitness gradient of a mutant *x*_*m*_ relative to a resident *x* in the case of decreasing exponential or sigmoïdes forms of the attack rate (top and bottom respectively), for a CO_2_ concentration worth 0.1, 15 and 50 µmol.L^-1^. Areas for which the fitness gradient of the mutant relative to the resident is negative are shown in white.

**Figure S2.**
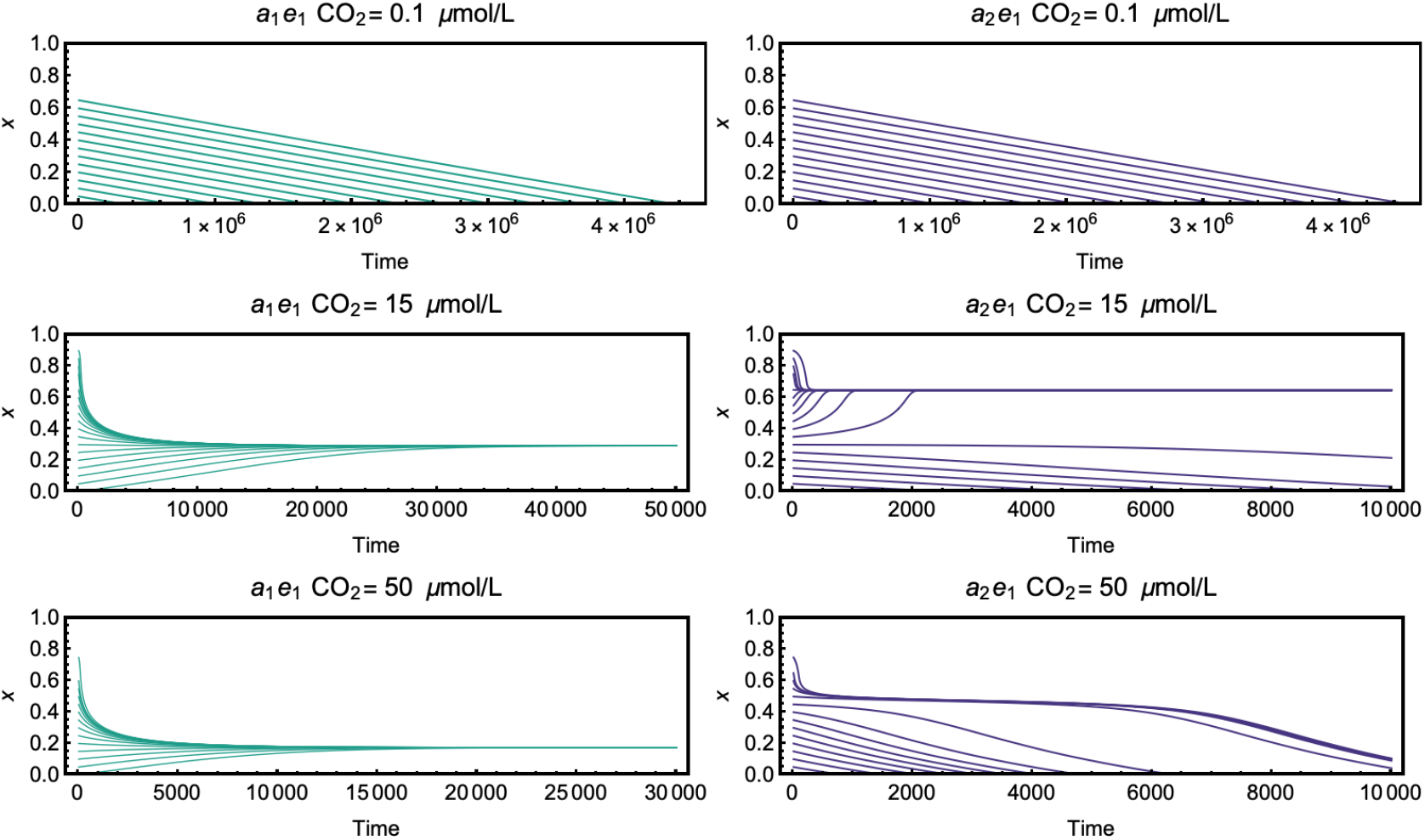
Temporal evolution of *x* for different initial phenotypes given two combinations of trade-off shapes and under CO_2_ concentrations of 0, 15 and 50 µmol.L^-1^. Integration of eq 18 has been completed using NDSolve of Mathematica. All other parameters are fixed for values shown Table 2.

**Figure S3.**
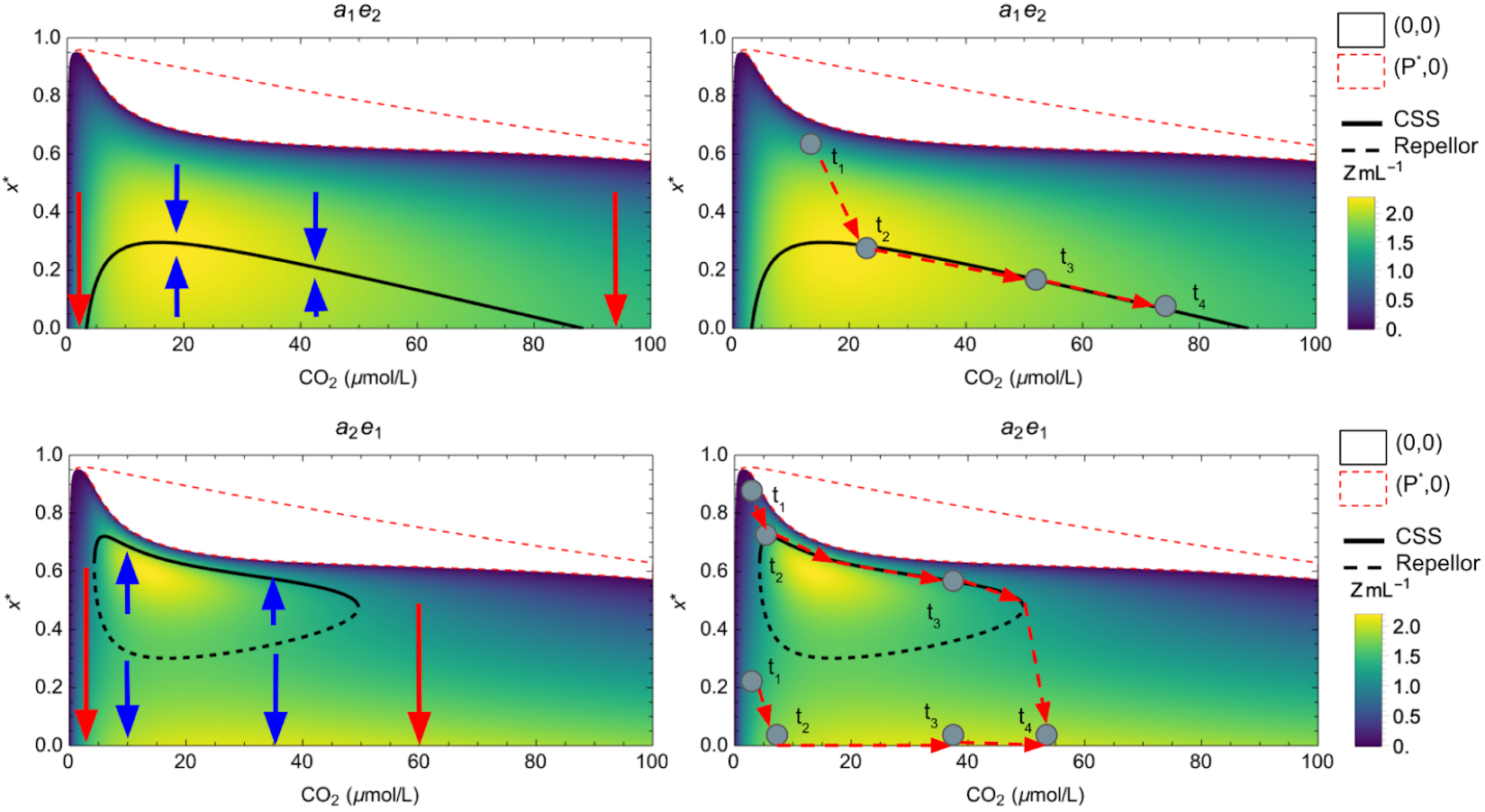
E_3_-diagram : selected strategy (continuous line : CSS, discontinuous line : Repellor) in the case of remaining tradeoff shapes combinations. Arrows : direction of evolution. A fictive scenario of the evolution of *x* over time (right panels) is represented for ease of understanding. Zooplankton densities are also represented. Light gray area : (*x, CO*_*2*_) space excluding only zooplankton. Dark gray area : (*x, CO*_*2*_) space where no species is able to survive. All other parameters are fixed at the values shown in Table 2.

**Figure S4.**
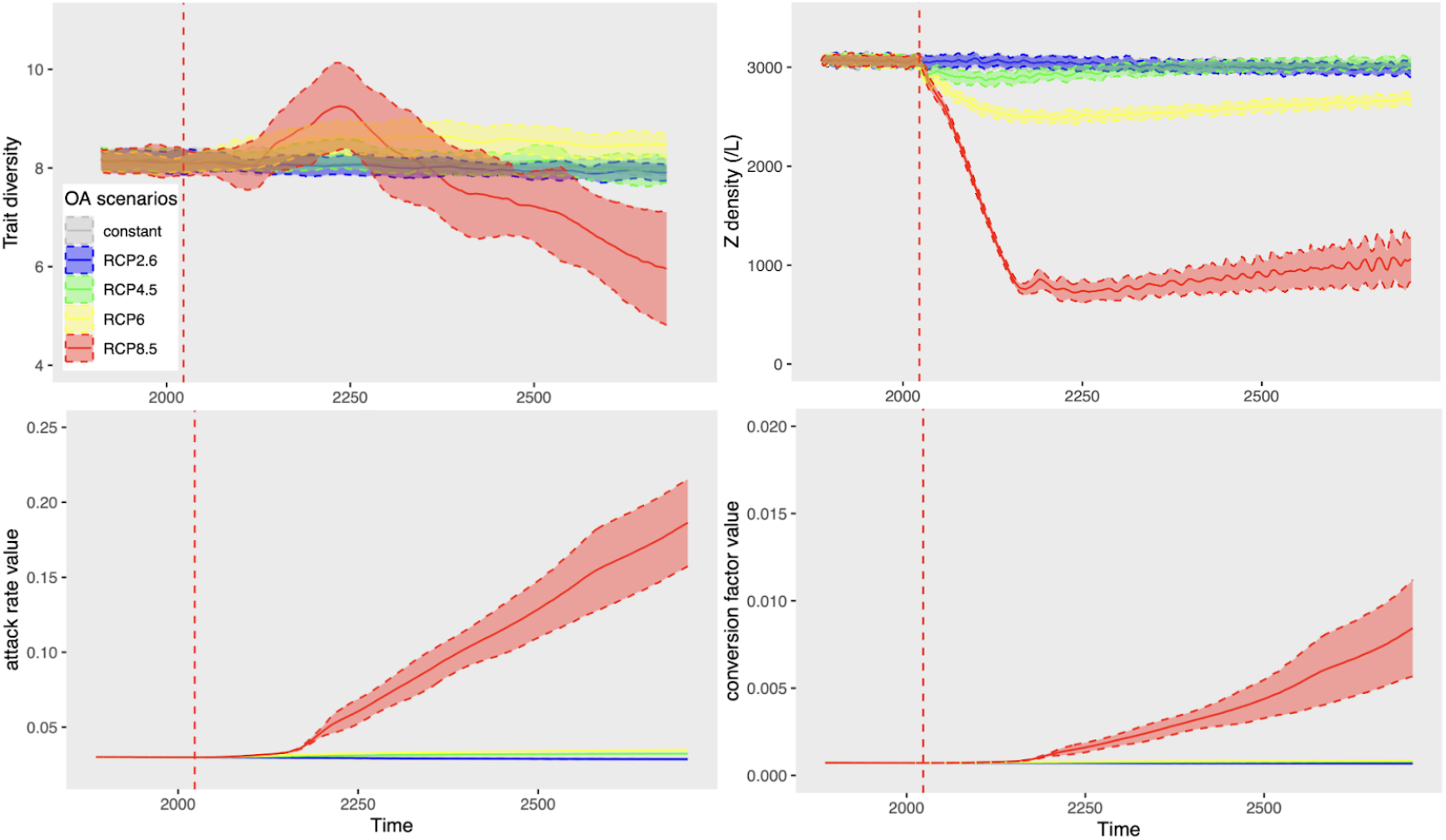
Impacts of RCP scenarios on coccolithophorids trait diversity, zooplankton densities, attack rates and conversion abilities. Full line : mean of 50 simulations for every RCP scenario. Dashed-line : standard deviation

## References

Anthony, K.R.N., Kline, D.I., Diaz-Pulido, G., Dove, S., Hoegh-Guldberg, O., 2008. Ocean acidification causes bleaching and productivity loss in coral reef builders. Proc. Natl. Acad. Sci. 105, 17442–17446. 10.1073/pnas.0804478105

Bach, L.T., Bauke, C., Meier, K.J.S., Riebesell, U., Schulz, K.G., 2012. Influence of changing carbonate chemistry on morphology and weight of coccoliths formed by &lt;i&gt;Emiliania huxleyi&lt;/i&gt; Biogeosciences 9, 3449–3463. 10.5194/bg-9-3449-2012

Bach, L.T., Riebesell, U., Gutowska, M.A., Federwisch, L., Schulz, K.G., 2015. A unifying concept of coccolithophore sensitivity to changing carbonate chemistry embedded in an ecological framework. Prog. Oceanogr. 135, 125–138. 10.1016/j.pocean.2015.04.012

Bach, L.T., Riebesell, U., Schulz, K.G., 2011. Distinguishing between the effects of ocean acidification and ocean carbonation in the coccolithophore Emiliania huxleyi. Limnol. Oceanogr. 56, 2040–2050. 10.4319/lo.2011.56.6.2040

Bijma, J., Pörtner, H.-O., Yesson, C., Rogers, A.D., 2013. Climate change and the oceans – What does the future hold? Mar. Pollut. Bull. 74, 495–505. 10.1016/j.marpolbul.2013.07.022

Brownlee, C., Taylor, A., 2004. Calcification in coccolithophores: A cellular perspective, in: Thierstein, H.R., Young, J.R. (Eds.), Coccolithophores. Springer Berlin Heidelberg, Berlin, Heidelberg, pp. 31–49. 10.1007/978-3-662-06278-4_2

Caldeira, K., Wickett, M.E., 2003. Anthropogenic carbon and ocean pH. Nature 425, 365–365. 10.1038/425365a

Cooper, R.N., Houghton, J.T., McCarthy, J.J., Metz, B., 2002. Climate Change 2001: The Scientific Basis. Foreign Aff. 81, 208. 10.2307/20033020

Cripps, G., Lindeque, P., Flynn, K.J., 2014. Have we been underestimating the effects of ocean acidification in zooplankton? Glob. Change Biol. 20, 3377–3385. 10.1111/gcb.12582

Dakos, V., Matthews, B., Hendry, A.P., Levine, J., Loeuille, N., Norberg, J., Nosil, P., Scheffer, M., De Meester, L., 2019. Ecosystem tipping points in an evolving world. Nat. Ecol. Evol. 3, 355–362. 10.1038/s41559-019-0797-2

De Mazancourt, C., Dieckmann, U., 2004. Trade-Off Geometries and Frequency-Dependent Selection. Am. Nat. 164, 765–778. 10.1086/424762

Denman, K.L., Brasseur, G., Chidthaisong, A., Ciais, P., Cox, P.M., Dickinson, R.E., Hauglustaine, D., Heinze, C., Holland, E., Jacob, D., Lohmann, U., Ramachandran, S., Archer, D., Arora, V., Austin, J., Baker, D., Berry, J.A., Betts, R., Bonan, G., Bousquet, P., Canadell, J., Christian, J., Clark, D.A., Dameris, M., Dentener, F., Easterling, D., Eyring, V., Feichter, J., Friedlingstein, P., Fung, I., Fuzzi, S., Gong, S., Gruber, N., Guenther, A., Gurney, K., Henderson-Sellers, A., House, J., Jones, A., Jones, C., Kärcher, B., Kawamiya, M., Lassey, K., Leck, C., Lee-Taylor, J., Malhi, Y., Masarie, K., McFiggans, G., Menon, S., Miller, J.B., Peylin, P., Pitman, A., Quaas, J., Raupach, M., Rayner, P., Rehder, G., Riebesell, U., Rödenbeck, C., Rotstayn, L., Roulet, N., Sabine, C., Schultz, M.G., Schulz, M., Schwartz, S.E., Steffen, W., Stevenson, D., Tian, Y., Trenberth, K.E., Noije, T.V., Wild, O., Zhang, T., Zhou, L., Boonpragob, K., Heimann, M., Molina, M., n.d. Couplings Between Changes in the Climate System and Biogeochemistry.

Dieckmann, U., Law, R., n.d. The dynamical theory of coevolution: a derivation from stochastic ecological processes.

Doebeli, M., Dieckmann, U., n.d. Evolutionary branching and sympatric speciation.

Doney, S.C., Fabry, V.J., Feely, R.A., Kleypas, J.A., 2009. Ocean Acidification: The Other CO _2_ Problem. Annu. Rev. Mar. Sci. 1, 169–192. 10.1146/annurev.marine.010908.163834

Dutkiewicz, S., Morris, J.J., Follows, M.J., Scott, J., Levitan, O., Dyhrman, S.T., Berman-Frank, I., 2015. Impact of ocean acidification on the structure of future phytoplankton communities. Nat. Clim. Change 5, 1002–1006. 10.1038/nclimate2722

Erwin, D., 2008. Macroevolution of ecosystem engineering, niche construction and diversity. Trends Ecol. Evol. 23, 304–310. 10.1016/j.tree.2008.01.013

Eshel, I., n.d. Evolutionary and Continnoas Stability.

Fabry, V.J., Seibel, B.A., Feely, R.A., Orr, J.C., 2008. Impacts of ocean acidification on marine fauna and ecosystem processes. ICES J. Mar. Sci. 65, 414–432. 10.1093/icesjms/fsn048

Feely, R.A., Sabine, C.L., Hernandez-Ayon, J.M., Ianson, D., Hales, B., 2008. Evidence for Upwelling of Corrosive “Acidified” Water onto the Continental Shelf. Science 320, 1490–1492. 10.1126/science.1155676

Feely, R.A., Sabine, C.L., Lee, K., Berelson, W., Kleypas, J., Fabry, V.J., Millero, F.J., 2004. Impact of Anthropogenic CO _2_ on the CaCO _3_ System in the Oceans. Science 305, 362–366. 10.1126/science.1097329

Ferriere, R., Legendre, S., 2013. Eco-evolutionary feedbacks, adaptive dynamics and evolutionary rescue theory. Philos. Trans. R. Soc. B Biol. Sci. 368, 20120081. 10.1098/rstb.2012.0081

Field, C.B., Behrenfeld, M.J., Randerson, J.T., Falkowski, P., 1998. Primary Production of the Biosphere: Integrating Terrestrial and Oceanic Components. Science 281, 237–240. 10.1126/science.281.5374.237

Gafar, N.A., Eyre, B.D., Schulz, K.G., 2018. A Conceptual Model for Projecting Coccolithophorid Growth, Calcification and Photosynthetic Carbon Fixation Rates in Response to Global Ocean Change. Front. Mar. Sci. 4, 433. 10.3389/fmars.2017.00433

Gattuso, J., 1998. Effect of calcium carbonate saturation of seawater on coral calcification. Glob. Planet. Change 18, 37–46. 10.1016/S0921-8181(98)00035-6

Geritz, S.A.H., Kisdi, É., Meszéna, G., Metz, J.A.J., 1998. Evolutionarily singular strategies and the adaptive growth and branching of the evolutionary tree. Evol. Ecol. 12, 35–57. 10.1023/A:1006554906681

Harvey, E.L., Bidle, K.D., Johnson, M.D., 2015. Consequences of strain variability and calcification in Emiliania huxleyi on microzooplankton grazing. J. Plankton Res. fbv081. 10.1093/plankt/fbv081

Haunost, M., Riebesell, U., D’Amore, F., Kelting, O., Bach, L.T., 2021. Influence of the Calcium Carbonate Shell of Coccolithophores on Ingestion and Growth of a Dinoflagellate Predator. Front. Mar. Sci. 8, 664269. 10.3389/fmars.2021.664269

Herms, D.A., Mattson, W.J., 1992. The Dilemma of Plants: To Grow or Defend. Q. Rev. Biol. 67, 283–335. 10.1086/417659

Hoegh-Guldberg, O., Poloczanska, E.S., Skirving, W., Dove, S., 2017. Coral Reef Ecosystems under Climate Change and Ocean Acidification. Front. Mar. Sci. 4, 158. 10.3389/fmars.2017.00158

Holligan, P.M., Robertson, J.E., 1996. Significance of ocean carbonate budgets for the global carbon cycle. Glob. Change Biol. 2, 85–95. 10.1111/j.1365-2486.1996.tb00053.x

Hoppe, C.J.M., Wolf, K.K.E., Schuback, N., Tortell, P.D., Rost, B., 2018. Compensation of ocean acidification effects in Arctic phytoplankton assemblages. Nat. Clim. Change 8, 529–533. 10.1038/s41558-018-0142-9

Intergouvernemental panel on climate change (Ed.), 2007. Climate change 2007: the physical science basis. Cambridge university press, Cambridge.

Jansen, H., Zeebe, R.E., Wolf-Gladrow, D.A., 2002. Modeling the dissolution of settling CaCO _3_ in the ocean. Glob. Biogeochem. Cycles 16. 10.1029/2000GB001279

Kleypas, J., Yates, K., 2009. Coral Reefs and Ocean Acidification. Oceanography 22, 108–117. 10.5670/oceanog.2009.101

Kolb, A., Strom, S., 2013. An inducible antipredatory defense in haploid cells of the marine microalga Emiliania huxleyi (Prymnesiophyceae). Limnol. Oceanogr. 58, 932–944. 10.4319/lo.2013.58.3.0932

Kroeker, K.J., Kordas, R.L., Crim, R., Hendriks, I.E., Ramajo, L., Singh, G.S., Duarte, C.M., Gattuso, J., 2013. Impacts of ocean acidification on marine organisms: quantifying sensitivities and interaction with warming. Glob. Change Biol. 19, 1884–1896. 10.1111/gcb.12179

Kroeker, K.J., Kordas, R.L., Crim, R.N., Singh, G.G., 2010. Meta-analysis reveals negative yet variable effects of ocean acidification on marine organisms. Ecol. Lett. 13, 1419–1434. 10.1111/j.1461-0248.2010.01518.x

Kuffner, I.B., Andersson, A.J., Jokiel, P.L., Rodgers, K.S., Mackenzie, F.T., 2008. Decreased abundance of crustose coralline algae due to ocean acidification. Nat. Geosci. 1, 114–117. 10.1038/ngeo100

Leibold, M.A., 1996. A Graphical Model of Keystone Predators in Food Webs: Trophic Regulation of Abundance, Incidence, and Diversity Patterns in Communities. Am. Nat. 147, 784–812. 10.1086/285879

Macarthur, R., Levins, R., 1967. The Limiting Similarity, Convergence, and Divergence of Coexisting Species. Am. Nat. 101, 377–385. 10.1086/282505

McNeil, B.I., Matear, R.J., n.d. Southern Ocean acidification: A tipping point at 450-ppm atmospheric CO2.

Meakin, N.G., Wyman, M., 2011. Rapid shifts in picoeukaryote community structure in response to ocean acidification. ISME J. 5, 1397–1405. 10.1038/ismej.2011.18

Metz, J.A.J., 1992. How ShouldWe Define’Fitness’for GeneralEcologicaSl cenarios? 7.

Meyer, J., Riebesell, U., 2015. Reviews and Syntheses: Responses of coccolithophores to ocean acidification: a meta-analysis. Biogeosciences 12, 1671–1682. 10.5194/bg-12-1671-2015

Millero, F.J., n.d. of the carbon dioxide system in the oceans.

Milliman, J.D., 1993. Production and accumulation of calcium carbonate in the ocean: Budget of a nonsteady state. Glob. Biogeochem. Cycles 7, 927–957. 10.1029/93GB02524

Monteiro, F.M., Bach, L.T., Brownlee, C., Bown, P., Rickaby, R.E.M., Poulton, A.J., Tyrrell, T., Beaufort, L., Dutkiewicz, S., Gibbs, S., Gutowska, M.A., Lee, R., Riebesell, U., Young, J., Ridgwell, A., 2016. Why marine phytoplankton calcify. Sci. Adv. 2, e1501822. 10.1126/sciadv.1501822

Orr, J.C., Kwiatkowski, L., Pörtner, H.-O., 2022. Arctic Ocean annual high in PCO2 could shift from winter to summer. Nature 610, 94–100. 10.1038/s41586-022-05205-y

Pandolfi, J.M., Connolly, S.R., Marshall, D.J., Cohen, A.L., 2011. Projecting Coral Reef Futures Under Global Warming and Ocean Acidification. Science 333, 418–422. 10.1126/science.1204794

Paul, A.J., Bach, L.T., 2020. Universal response pattern of phytoplankton growth rates to increasing CO _2_. New Phytol. 228, 1710–1716. 10.1111/nph.16806

Poulton, A.J., Adey, T.R., Balch, W.M., Holligan, P.M., 2007. Relating coccolithophore calcification rates to phytoplankton community dynamics: Regional differences and implications for carbon export. Deep Sea Res. Part II Top. Stud. Oceanogr. 54, 538–557. 10.1016/j.dsr2.2006.12.003

Quéré, C.L., Harrison, S.P., Colin Prentice, I., Buitenhuis, E.T., Aumont, O., Bopp, L., Claustre, H., Cotrim Da Cunha, L., Geider, R., Giraud, X., Klaas, C., Kohfeld, K.E., Legendre, L., Manizza, M., Platt, T., Rivkin, R.B., Sathyendranath, S., Uitz, J., Watson, A.J., Wolf-Gladrow, D., 2005. Ecosystem dynamics based on plankton functional types for global ocean biogeochemistry models. Glob. Change Biol. 11, 2016–2040. 10.1111/j.1365-2486.2005.1004.x

Riahi, K., Rao, S., Krey, V., Cho, C., Chirkov, V., Fischer, G., Kindermann, G., Nakicenovic, N., Rafaj, P., 2011. RCP 8.5—A scenario of comparatively high greenhouse gas emissions. Clim. Change 109, 33–57. 10.1007/s10584-011-0149-y

Riebesell, U., Wolf-Gladrow, D.A., Smetacek, V., 1993. Carbon dioxide limitation of marine phytoplankton growth rates. Nature 361, 249–251. 10.1038/361249a0

Riebesell, U., Zondervan, I., Zeebe, R.E., 2000. Reduced calci®cation of marine plankton in response to increased atmospheric CO2 407.

Sabine, C.L., Feely, R.A., Gruber, N., Key, R.M., Lee, K., Bullister, J.L., Wanninkhof, R., Wong, C.S., Wallace, D.W.R., Tilbrook, B., Millero, F.J., Peng, T.-H., Kozyr, A., Ono, T., Rios, A.F., 2004. The Oceanic Sink for Anthropogenic CO _2_. Science 305, 367–371. 10.1126/science.1097403

Scheffer, M., Carpenter, S.R., 2003. Catastrophic regime shifts in ecosystems: linking theory to observation. Trends Ecol. Evol. 18, 648–656. 10.1016/j.tree.2003.09.002

Schlüter, L., Lohbeck, K.T., Gröger, J.P., Riebesell, U., Reusch, T.B.H., 2016. Long-term dynamics of adaptive evolution in a globally important phytoplankton species to ocean acidification. Sci. Adv. 2, e1501660. 10.1126/sciadv.1501660

Schlüter, L., Lohbeck, K.T., Gutowska, M.A., Gröger, J.P., Riebesell, U., Reusch, T.B.H., 2014. Adaptation of a globally important coccolithophore to ocean warming and acidification. Nat. Clim. Change 4, 1024–1030. 10.1038/nclimate2379

Sett, S., Bach, L.T., Schulz, K.G., Koch-Klavsen, S., Lebrato, M., Riebesell, U., 2014. Temperature Modulates Coccolithophorid Sensitivity of Growth, Photosynthesis and Calcification to Increasing Seawater pCO2. PLoS ONE 9, e88308. 10.1371/journal.pone.0088308

Strom, S.L., Bright, K.J., Fredrickson, K.A., Cooney, E.C., 2018. Phytoplankton defenses: Do Emiliania huxleyi coccoliths protect against microzooplankton predators? Limnol. Oceanogr. 63, 617–627. 10.1002/lno.10655

Taucher, J., Haunost, M., Boxhammer, T., Bach, L.T., Algueró-Muñiz, M., Riebesell, U., 2017. Influence of ocean acidification on plankton community structure during a winter-to-summer succession: An imaging approach indicates that copepods can benefit from elevated CO2 via indirect food web effects. PLOS ONE 12, e0169737. 10.1371/journal.pone.0169737

Westbroek, P., Young, J.R., Linschooten, K., 1989. Coccolith Production (Biomineralization) in the Marine Alga Emiliania huxleyi. J. Protozool. 36, 368–373. 10.1111/j.1550-7408.1989.tb05528.x

Young, J.R., Andruleit, H., Probert, I., 2009. COCCOLITH FUNCTION AND MORPHOGENESIS: INSIGHTS FROM APPENDAGE-BEARING COCCOLITHOPHORES OF THE FAMILY SYRACOSPHAERACEAE (HAPTOPHYTA) ^1^. J. Phycol. 45, 213–226. 10.1111/j.1529-8817.2008.00643.x

